# Astroglia proliferate upon biogenesis of tunneling nanotubes and clearance of α-synuclein toxicities

**DOI:** 10.1101/2023.08.24.554645

**Authors:** Abinaya Raghavan, Rachana Kashyap, P Sreedevi, Sneha Jos, Suchana Chatterjee, Ann Alex, Michelle Ninochka D’Souza, Mridhula Giridharan, Ravi Muddashetty, Ravi Manjithaya, Sivaraman Padavattan, Sangeeta Nath

## Abstract

Astrocytic cells are a subtype of glial cells that engulf pathogenic aggregates derived from degenerative neurons to facilitate its degradation. Here, we show that exposure to α-SYN protofibrils caused a transient increase in biogenesis of tunneling nanotubes (TNTs) in primary astrocytes and astrocyte-origin cancer cell-lines (U-87 MG, U251). Biogenesis of nascent TNTs corresponds to α-SYN protofibril-induced organelle toxicities, increased reactive oxygen species (ROS), and oxidative stress-induced premature cellular senescence. These TNTs mediate cell-to-cell transfer of α-SYN protofibrils, toxic lysosomes and mitochondria. Biogenesis of TNTs precedes clearance of α-SYN-induced organelle toxicities, cellular ROS levels and reversal of cellular senescence. Consequences of cellular clearance results in enhanced cell proliferation. Further, we have shown α-SYN-induced senescence promotes transient localization of focal adhesion kinase (FAK) in the nucleus. FAK mediated regulation of Rho-associated kinases may have a role in the biogenesis of TNTs, successively proliferation. Our study emphasizes that TNT biogenesis may have a potential role in the clearance of α-SYN toxicities and reversal of stress-induced cellular senescence, consequences of which cause enhanced proliferation in the post-recovered astroglia cells.

**Highlights:**

- α-SYN protofibrils treated astroglia cells proliferate upon transient biogenesis of TNTs.
- Transient TNT biogenesis precedes clearance of α-SYN toxicities and reversal of senescence.
- Stress-induced senescence results in nuclear localization of FAK and ROCK mediated TNT biogenesis.
- The rescued cells enhance proliferation through ROCK mediated ERK1/2 and NFκB signalling cascades.

**Synopsis:** Graphical Abstract:
α-SYN protofibrils-induced biogenesis of tunneling nanotubes (TNTs) aids to enhance cellular clearance of toxic burdens as a cellular survival strategy. α-SYN protofibrils treated toxic senescence cells regulate FAK mediated modulation of ROCK signalling cascades to promote TNT biogenesis and rescue the cellular toxicities. The rescued cells eventually enhance cell proliferation.

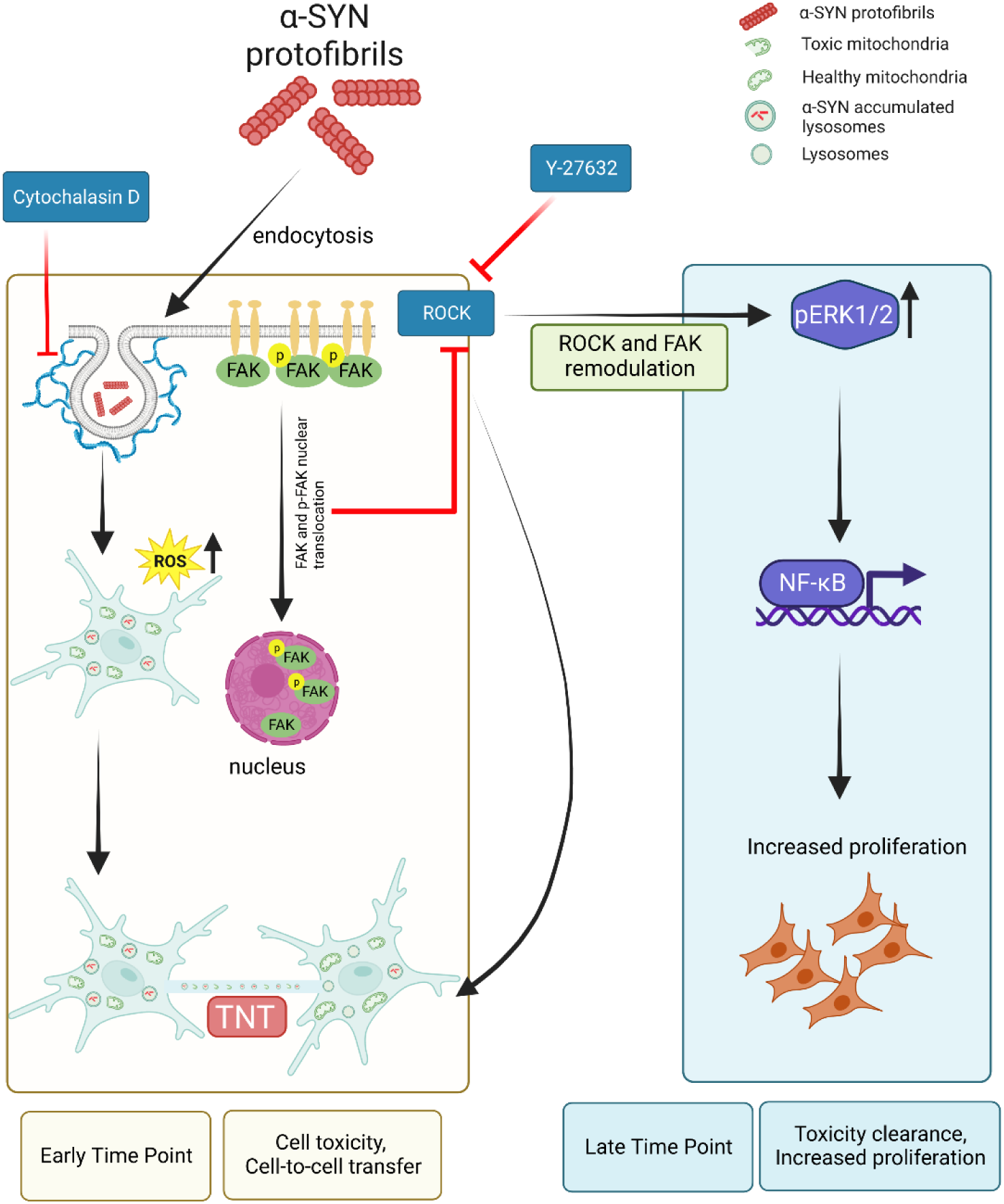

## Introduction

Parkinson’s disease (PD) is characterized by the loss of dopaminergic neurons in the *substantia nigra* (SN) due to cytoplasmic accumulation of Lewy bodies or Lewy neurites which mainly consists of α-synuclein (α-SYN) protein (Forno 1996, Kalia and Lang 2015). Along with the accumulated cytoplasmic α-SYN, extracellular α-SYN also plays a significant role in neurodegeneration, progressive intercellular spreading of pathology and neuroinflammation (Lee, Bae et al. 2014, Yamada and Iwatsubo 2018). Studies have shown that elevation of neuronal activity and various stress conditions increase extracellular release of α-SYN (Fortin, Nemani et al. 2005, Yamada and Iwatsubo 2018). Pathogenic aggregates of α-SYN can release from degenerating neurons, which can be taken up by surrounding neurons and glial cells. Glial cells normally express low levels of α-SYN, however, at the advanced stage of PD, α-SYN aggregates in astrocytes and glial synucleinopathies are often detected (Krejciova, Carlson et al. 2019). Additionally, gliosis is a typical pathological feature of neurodegenerative diseases. Sustained activation and fibrous proliferation of glial-cells, mainly astrocytes and microglia, are central features of dopaminergic neurodegeneration in PD (MacMahon Copas, McComish et al. 2021). Neuronal activity modulates cellular cross-talks and intercellular communication between neurons and the surrounding neuroglial-cells (Gibson, Purger et al. 2014, Venkatesh,Johung et al. 2015, Venkatesh, Morishita et al. 2019).

Recent studies have shown that mode of intercellular transfer of neurodegenerative proteins between glial cells facilitate the clearing of neurodegenerative aggregates, such as α-SYN and amyloid-β (Aβ) (Rostami, Holmqvist et al. 2017, Rostami, Mothes et al. 2021, Scheiblich, Dansokho et al. 2021). The spreading of neurodegenerative proteins through an intercellular mode of transfer and its role in pathology progression have widely been studied in several model systems (Victoria and Zurzolo 2017, Neupane, De Cecco et al. 2022). Exosomes, unconventional secretion and recently, direct cell-to-cell transfer via membrane nanotubes or tunneling nanotubes (TNTs) have been demonstrated by several studies as modes of intercellular transfer (Neupane, De Cecco et al. 2022).

Recent discoveries of TNTs have opened up the possibility of direct long-range cell-to-cell communication (Rustom, Saffrich et al. 2004). TNTs have been shown to be open-ended, and thin (diameter around 50-700 nm) while having an intercellular membrane-actin continuity (long upto 300 µm) between distant cells. TNTs are reported to be a conduit for direct transfer of organelles (Rustom, Saffrich et al. 2004), neurodegeneration-associated protein aggregates (Victoria and Zurzolo 2017, Ramirez-Jarquin, Sharma et al. 2022), viruses (Jansens, Tishchenko et al. 2020) and RNA (Haimovich, Ecker et al. 2017) between cells. Neurodegenerative aggregate-induced endo-lysosomal toxicities and mitochondrial stress promote biogenesis of TNTs (Victoria and Zurzolo 2017, Raghavan, Rao et al. 2021). Propagation of pathogenic aggregates *via* TNTs have also shown to aid in the progression of neurodegeneration (Rostami, Holmqvist et al. 2017, Victoria and Zurzolo 2017, Dilna, Deepak et al. 2021). In this context, several studies have shown that α-SYN aggregates in lysosomes transfer from neuron to neuron *via* TNTs in PD (Abounit, Bousset et al. 2016, Victoria and Zurzolo 2017, Dilsizoglu Senol, Samarani et al. 2021). On the other hand, TNTs mediate crosstalk between astrocytes, and microglia to degrade toxic α-SYN through cell-to-cell transfer of aggregates and help to reduce ROS accumulation and oxidative stress (Rostami, Holmqvist et al. 2017, Scheiblich, Dansokho et al. 2021). Oxidative stress and ROS promote formation of TNTs and cell-to-cell transfer (Wang, Cui et al. 2011, Desir, Dickson et al. 2016).

ROS induced by the accumulation of neurotoxic proteins in neurons can induce cell senescence and that can aggravate neurodegeneration (Zizhen Si 2021, Yoon, You et al. 2022). Cellular senescence is a biologically homeostatic phenomenon that is defined by a persistent degree of cell cycle inhibition and cellular ageing. Cell cycle arrest in senescence cells is believed to be irreversible, however, studies have shown that stress-induced premature senescence that is induced by DNA damage-mediated amplification of the p53/p21 signalling pathway can be reversed by modulating the levels of p53 and p21 expressions (Macip, Igarashi et al. 2002, Beausejour, Krtolica et al. 2003).

Senescent cells are more prevalent in brain tissues and exert roles in brain ageing and neurodegeneration (Martinez-Cue and Rueda 2020). Brain cells including astrocytes, microglia, oligodendrocytes, and epithelial cells undergo cellular senescence under oxidative stress, which aggravates neurodegeneration (Schousboe, Bak et al. 2013). On the contrary, glial cells, especially astrocytes and microglia, play a critical role in protecting neurons by clearing toxic accumulations of neurodegenerative proteins from neurons and brains (Rostami, Holmqvist et al. 2017, Scheiblich, Dansokho et al. 2021). α-SYN induced toxicities can produce cellular senescence in response to DNA damage (Yoon, You et al. 2022). Thus, investigation into the mechanisms of TNT biogenesis and role of TNTs in facilitating clearance of toxic α-SYN protofibrils from brain to improve cell survival is needed.

In this study, we treated primary astrocytes, and astrocyte origin U-87 MG, U251 cancer cells with toxic α-SYN protofibrils and observed that the endo-lysosomal accumulation of α-SYN protofibrils induce mitochondrial toxicities, and thereby, trigger the biogenesis of membrane nanotubes or TNTs in a transient manner. Biogenesis of TNTs precedes clearance of α-SYN-induced organelle toxicities and cellular stress induced premature senescence. Further, we observed ROCK signalling modulates biogenesis of TNTs, and successively enhances cell proliferation in the post-recovered astroglia cells. Therefore, this study emphasizes the potential role of TNTs in facilitating cellular clearance in pathological stress conditions which maintains the survival of astroglial cell, probably by enhancing cell proliferation.

## Results

### α-SYN protofibrils induce biogenesis of transient TNTs

The contributions of extracellular α-SYN protofibrils in pathological spreading of PD in neuronal cell culture models are well studied (Dieriks, Park et al. 2017). Recent studies have shown that aggregates of α-SYN can modulate cellular cross-talks between neurons and the surrounding neuroglial cells (Chakraborty, Nonaka et al. 2023), which plays important role to get rid of toxic loads of aggregates (Rostami, Holmqvist et al. 2017, Scheiblich, Dansokho et al. 2021). However, the mechanism of establishment of intercellular communications and their fate are yet to be understood. Therefore, we looked into the primary astrocytes and astrocyte origin cancer cell lines (U-87 MG and U251) to understand the possible fate of the astroglia cells over time on α-SYN protofibril treatment.

To understand this, purified α-SYN was converted into protofibrils that were either unlabelled or fluorescently (TMR) labelled and characterized using transmission electron microscopy (Fig 1A). Interestingly, after treatment with 1µM α-SYN protofibrils at 3h, 6h, 12h and 24h, we observed formation of TNT-like membrane networks between primary astrocytes (Fig 1B), U-87 MG (Fig 1C, Fig EV1A) and U251 cells (Fig EV1B) for a transient time period at early hours. Time points were chosen quantifying optimum biogenesis of membrane nanotubes after treatment with 1µM α-SYN protofibrils over 24 h (Fig EV1C). Quantification of TNT-like structures were performed on confocal images of phalloidin-stained astrocytes (Fig 1B). DiD dye-stained membranes were imaged using fluorescence microscope to detect abundance of stained TNT-like membrane conduits in live cells (Fig 1C). DIC (Differential Interference Contrast) images of U-87 MG (Fig EV1A) and U251 (Fig EV1B) cells were also taken to detect TNT-like structures. Quantification showed increase in number of hovering thin TNT-like structures per cell compared to controls in the primary astrocytes (Fig 1D). DiD images of U-87 MG also revealed transient increase in the biogenesis of thin (diameters < 1 µm), cell-to-cell membrane connections at early time points (3h and 6h) (Fig 1E-F). Similarly, quantification of DIC images of U-87 MG cells (Fig EV1D) and U251 (Fig EV1E) showed enhanced transient biogenesis of TNTs at early time points compared to their respective controls and later time points.

**Figure 1:**
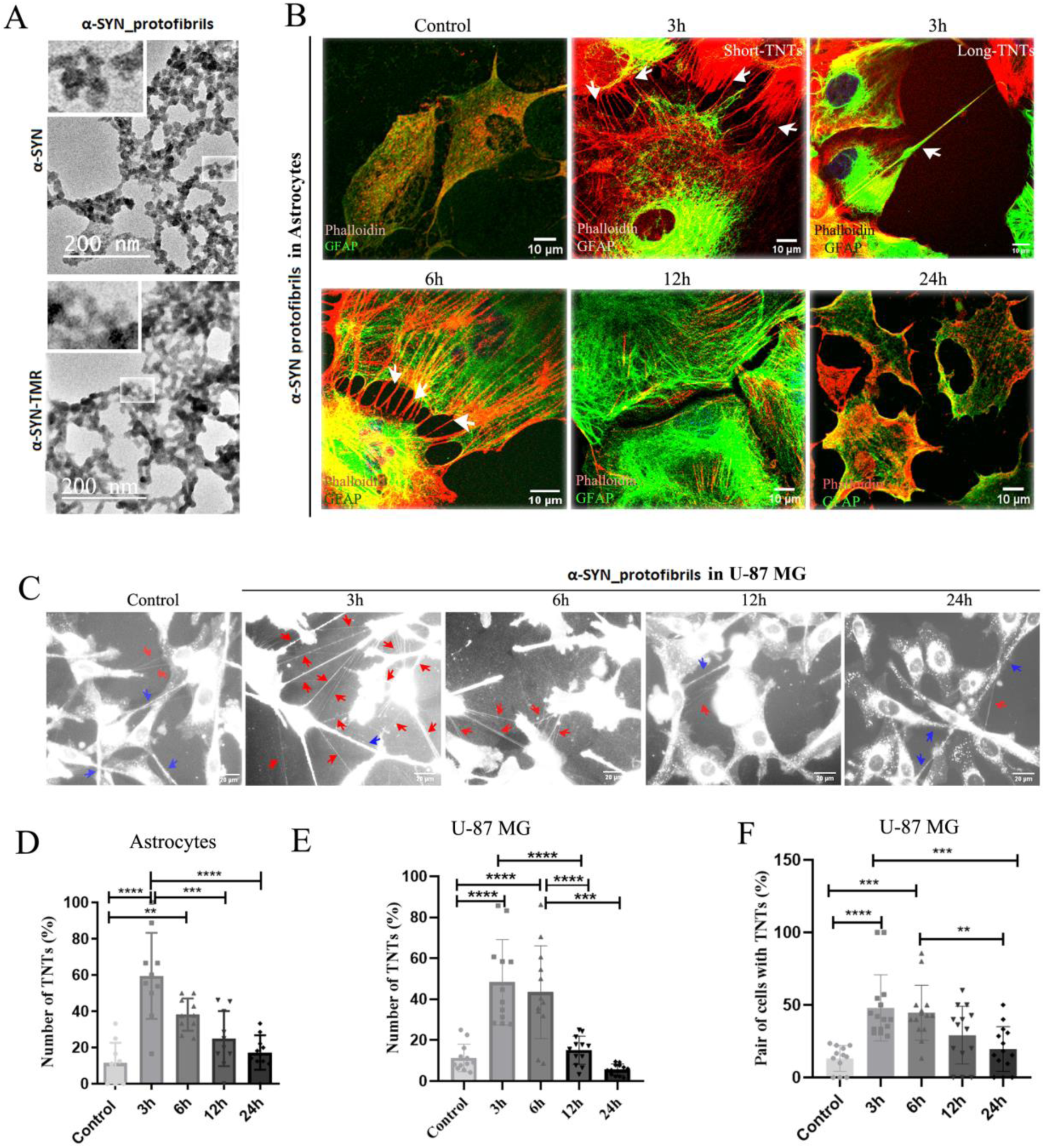
Characterization of α-SYN protofibrils and their effect on TNTs formation in astroglia cells. A) TEM (transmission electron microscope) images of α-SYN protofibrils unlabelled and TMR fluorescent dye labelled respectively. Magnified structures of protofibrils are shown in the zoomed panel. B) Primary astrocytes treated with 1µM α-SYN protofibrils for 3h, 6h, 12h and 24h, stained for actin (red) and GFAP (green), white arrows indicate formation of shorter and longer TNTs at 3h and 6h. C) DiD (membrane dye) stained U-87 MG cells showing transient increase of TNT-like structures (red arrows) in the cells treated with α-SYN protofibrils (1 μM) for 3 -6 h, compared to the respective controls and the cells treated for 12 h and 24 h. Blue arrows indicate relatively thicker tumour microtubes (TMs) like connections. D) Percentage of TNTs from primary astrocytes were quantified by counting the numbers normalized with the number of cells. E, F) Quantification of percentage of TNT-like structures and pairs of cells connected by them upon α-SYN protofibrils treatment in U-87 MG cells live stained with DiD. Quantifications are done from 10-15 image frames of a set and each image frame have 10-20 cells. Scale bars are denoted on the images. Data are expressed as mean ± SD, *** p ≤ 0.001. Statistics were analysed using two-way ANOVA. n=3.

### α-SYN protofibrils induce open-ended TNTs and cell-to-cell transfer

TNTs are actin rich, nano-sized in diameters and primarily defined as open-ended membrane tubes (Abounit, Delage et al. 2015). TNTs are distinguished from filopodia and neurites by their distinct characteristic to hover between two distant cells (Valappil, Raghavan et al. 2022). Actin-binding phalloidin and GFAP-stained, 3D-reconstructed confocal images show actin-positive thin TNTs that hover between two cells in primary astrocytes (Fig 2A). Astrocyte-origin cancer cell lines (U-87 MG and U251) are known to form tumour microtubes (TMs), close ended thicker membrane networks (diameters >1 µm) between cells and electrically coupled *via* gap junctions (Osswald, Jung et al. 2015). To distinguish TNTs and TMs, confocal z-stack images were obtained after immunocytochemical staining of cells using β-tubulin antibody and phalloidin dye. Thin actin-positive, hovering TNTs (Fig 2B) and thicker TMs (Fig 2C) are identified in U-87 MG cells. TMs are positive for both actin and β-tubulin. Studies have indicated that astrocytoma/glioblastoma cells express lower levels of neuronal marker β-tubulin and are stained faintly (Abbassi, Recasens et al. 2019). Despite that, β-tubulin staining is distinctly detected in thicker TMs and absent in TNTs (Fig EV2A and B). DIC images clearly showed the nano-sized diameters of TNTs and thicker TMs (Fig 2A-C-upper panels). Reconstructed 3D volume images of confocal z-stacks, show actin-positive, thin TNTs hover between two cells and are not attached to the substratum, whereas the tubulin-positive thicker TMs are present on the substratum (Fig 2A-C-lower panels).

**Figure 2:**
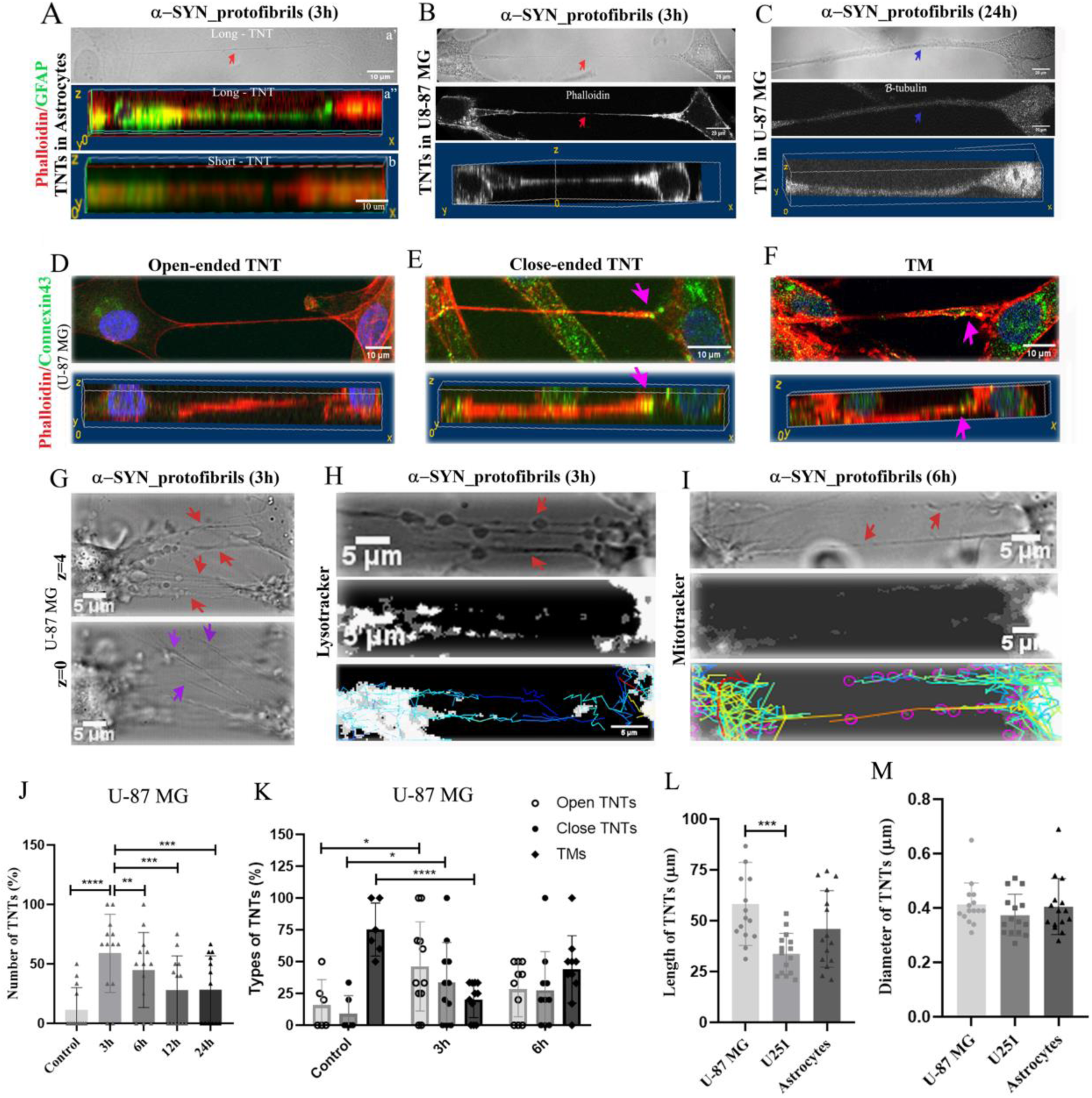
Characterization of TNTs formed upon treatment with α-SYN protofibrils in astroglia cells. A) Red arrow in the DIC image (a’) indicates TNT formed between two astrocytes on α-SYN protofibrils treatment. Bottom panel shows 3D volume view of Phalloidin and GFAP positive astrocytes connected by both long (a’’) and short TNTs (b). B, C) DIC images of U-87 MG cells show TNTs (B) and TMs (C). The cells were stained with phalloidin and β-tubulin, both bottom panels show 3D volume views of TNT as actin positive hovering structure (B) and tubulin positive TM at the surface (C). Red arrows indicate actin positive TNTs and blue arrows indicate β-tubulin positive tumour microtubes (TM) respectively. D, E) Characterization of open (D) and close (E) ended TNTs based on Phalloidin and connexin43 staining at the tip of the TNT in U-87 MG cells. Pink arrows indicate close-ended TNT. Bottom panel shows 3D volume views of the same. F) Closed ended TM characterized by the presence of connexin43 staining, indicated by pink arrows. Bottom panel shows 3D volume views of the TM at the surface. G) TNT-like structures (red arrows) are detectable in DIC images as focused structures at z=4. At z=0 filopodia like extensions (magenta arrows) on substratum are at focus. H, I) U87MG cells treated with 1µM α-SYN protofibrils for 3h and 6h, stained with lysotracker and mitotracker respectively. Red arrows indicate lysotracker and mitotracker through TNTs at 3h and 6h. Movement of lysotracker (H) and mitotracker (I) positive vesicles through TNTs were tracked using the trackmate plug-in of Fiji. J) Quantification of percentage of TNTs and K) percentage of open-ended TNTs, close-ended TNTs and TMs from the confocal Z-stack images of U-87 MG cells. Quantification of length (L) and diameter (M) of TNTs formed by U-87 MG cells, U251 cells and astrocytes. Quantifications are done from 10-15 image frames of a set and each image frame have 10-20 cells. Scale bars are denoted on the images. Data are expressed as mean ± SD, *** p ≤ 0.001. Statistics were analysed using two-way ANOVA (2J and 2K) and one-way ANOVA (2L and 2M). n=3.

Several studies have shown that TNTs are open-ended membrane channels, and can transfer organelles directly from one cell to another (Rustom, Saffrich et al. 2004, Abounit, Bousset et al. 2016). However, few studies have referred close ended membrane continuities that are not involved in cell-to-cell transfer of organelles and electrically coupled *via* gap junctions at the end point as TNTs (Sowinski, Jolly et al. 2008, Wang, Veruki et al. 2010). Therefore, TNTs were immunostained using gap junction protein connexin43, to identify open and close-ended TNTs. Images captured populations of both connexin43 negative (Fig 2D) and positive (Fig 2E) at the end point of TNTs. Non-hovering thicker TMs are connexin43 positive structures (Fig 2F).

We further functionally characterized TNTs based on their ability to transfer organelles like lysosomes and mitochondria. Time-lapse and z-stack images were taken using confocal microscopy (using both fluorescence channels and DIC channel) in live U-87 MG cells, stained with lysotracker and mitotracker, post treatment with 1µM α-SYN protofibrils (3h, 6h, 12h and 24h). Time-lapse videos in DIC microscopy images, revealed transient increase in the biogenesis of thin cell-to-cell membrane connections and unidirectional movements of organelles *via* the membrane nanotubes at early time points (3h and 6h) (Movies 1-2). Time-lapse videos captured cell-to-cell movement of α-SYN-TMR accumulated lysosomes and mitochondria through long (20 µm to 120 µm), thin TNTs after 3h and 6 h of treatments respectively (Movies 3 and 4). We validated from z-stack images, that the thin nano-sized membrane tubes or TNTs are hovering structures connecting between two distant cells. The DIC images show thin membrane nanotubes (red arrows) between cells that are clearly visible at a higher plane of z=4. The surface, at z=0, filopodia like membrane extensions (violet arrows) are visible (Fig 2G) which are unlike the TNTs. We tracked the movements of lysotracker (Fig 2H) and mitotracker-labelled (Fig 2I) organelles along the TNTs (identified from z-stacks) using TrackMate plug-in available in FIJI and we observed that both the organelles travel unidirectionally from one cell towards another, with an average speed of 0.062 ± 0.003 µm/sec. Further, we captured time-lapse videos to detect direct cell-to-cell transfer of α-SYN-TMR accumulated lysotracker and mitochondria *via* open-ended TNTs between U-87 MG cells after 3h of treatments (Movies 5 and 6). Our results reveal the formation of open-ended functional TNTs and cell-to-cell transfer through them at 3h and 6h post treatment with α-SYN.

We re-established transient increase in the biogenesis of thin (diameters < 1 µm), hovering TNTs upon 1µM α-SYN protofibrils treatment at early hours (3h and 6h) by quantifying from the 3D reconstruction of confocal z-stack images (Fig 2J). Though control cells contain higher percentage of TMs, we observed transient increase of both open and close-ended TNTs, based on connexin43 staining after 3h and 6h post treatment with α-SYN (Fig 2K). Length and diameters of TNTs were measured using the confocal z-stack images in all the cell types, primary astrocytes, U-87 MG and U251 cells. The careful measurements of z-stack images detected that the diameter of TNTs range between 1-3 pixels, that is around 220 nm to 660 nm and length varies between 20 µm – 100 µm across all the cell types (Fig 2L and M).

### Transient lysosomal toxicities in astroglia cells upon treatment with α-SYN protofibrils

Internalization of extracellularly applied α-SYN protofibrils into endo-lysosomal pathways induces transient accumulations and toxicities (3h and 6h) in primary astrocytes (Fig 3A) and U-87 MG cells (Fig 3B and C). α-SYN and LAMP1 were co-immunostained in primary astrocytes (Fig 3A) and U-87 MG cells treated with TMR-labelled α-SYN were stained with lysotracker (Fig 3B). Accumulations of α-SYN protofibrils in endo-lysosomes and resulting toxicities were studied following enlarged and distorted morphology of α-SYN accumulated endo-lysosomal vesicles at 3h and 6h (Fig 3A and B). Extent of cathepsin leakage from LAMP2-positive vesicles was also observed to follow lysosomal toxicities (Fig 3C). α-SYN protofibrils were internalized in primary astrocytes and in substantial amounts it colocalized to LAMP1-immunostained positive vesicles (analysed using Coloc 2 plug-in in FIJI). Maximum colocalizations were observed at 3h of treatment which over-time gradually decreases (Fig 3D). Internalization of protofibrils to lysosomal pathway caused transient accumulation and toxicities, resulting in larger sized LAMP1 positive vesicles transiently detectable at 3h and 6h after α-SYN protofibrils treatment in astrocytes (Fig 3E). Quantification of red puncta of α-SYN-TMR in terms of total area/content and size of each punctum, show that endo-lysosomal machineries in U-87 MG cells gradually clear up toxic protofibrils (Fig 3F). The analysis of cathepsin leakage in LAMP2-positive lysosomal vesicles indicated no significant increase of cathepsin-D to the cytosol (Fig 3C). However, we observed that the larger cathepsin-D and LAMP2-positive vesicles after 3h of protofibril treatments (Fig 3G). Morphology of LAMP2-positive lysosomes at 3h is elongated and rough in comparison to control, 6h, 12h and 24h treated cells (magnified images highlighted at upper-left corner of Fig 3C). The time of transient lysosomal toxicity coincides with the transient increase of TNT biogenesis (Fig 1B-F). Altogether the results suggest that transfer of organelles *via* TNTs probably aids the cells to cope with protofibril-induced lysosomal toxicities.

**Figure 3:**
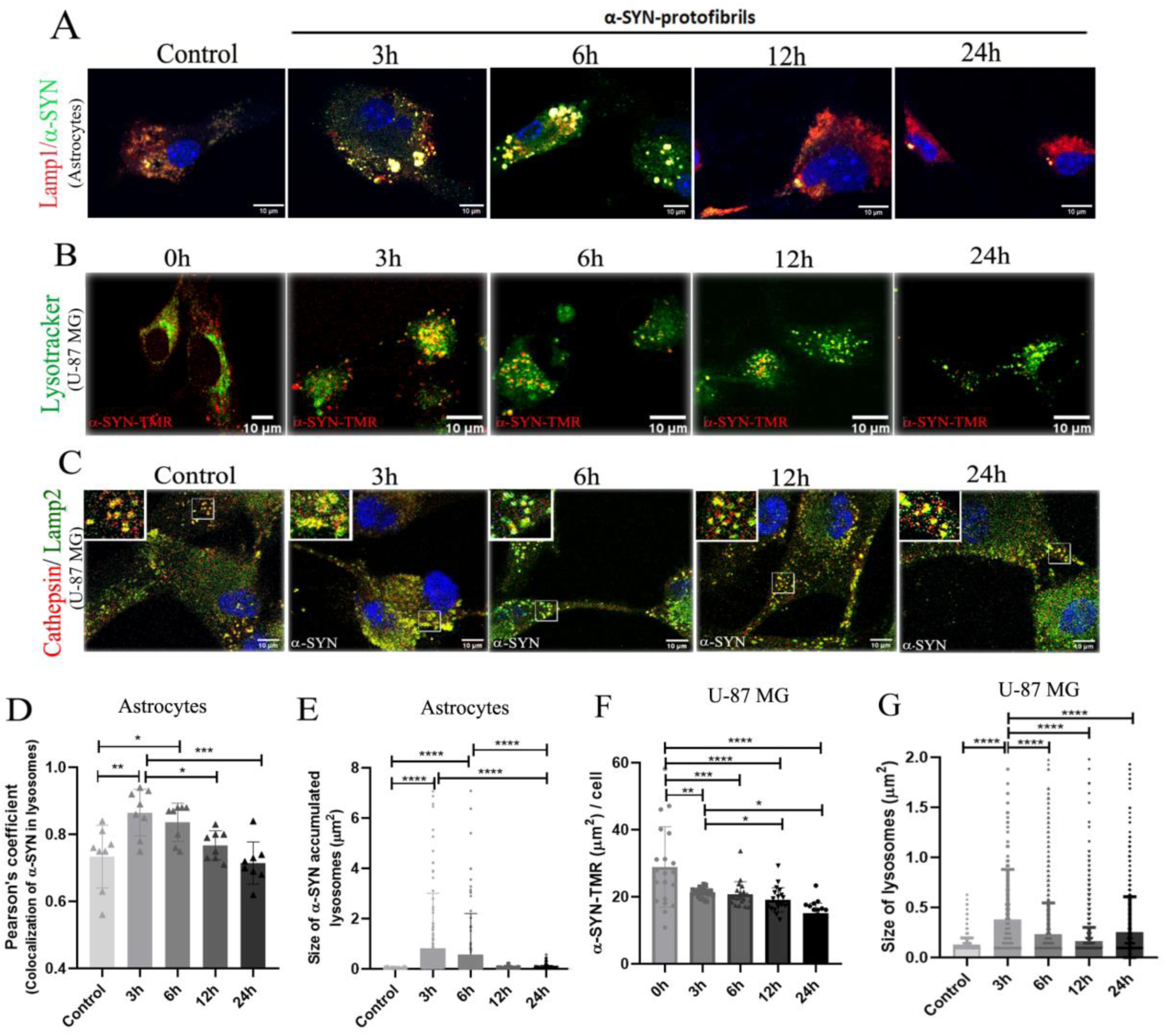
Effect of α-SYN protofibrils on lysosomes in U-87 MG cells and astrocytes. A) Mouse primary astrocytes treated with 1µM α-SYN protofibrils for 3h, 6h, 12h and 24h. α-SYN protofibrils (green) in lamp1 positive vesicles (red). At 3h and 6h larger sizes of lamp1 positive vesicles colocalized with α-SYN (yellow) are visible. B) U-87 MG cells treated with α-SYN-TMR protofibrils (red) in lysotracker positive vesicles (green). C) U-87 MG cells treated with α-SYN protofibrils (1µM) were stained using Cathepsin D (red) and Lamp2 (green). At 3h larger sizes of lamp2 positive vesicles colocalized with cathepsin-D were observed, compared to the other time points (magnified images highlighted at upper-left corner). D) Quantification of co-localization of α-SYN protofibrils in lamp1 positive lysosomes and (E) Size of α-SYN accumulated in lysosomes in astrocytes. F) Quantification of α-SYN accumulation per cell and G) Lysosome size in U-87 MG cells. Quantifications are done analysing randomly selected 10-15 cells from different image frames of an experimental set. Scale bars are denoted on the images. Data are expressed as mean ± SD, *** p ≤ 0.001. Statistics were analysed using two-way ANOVA. n=3.

### Transient mitochondrial toxicities in astroglia cells upon treatment with α-SYN protofibrils

Similar to transient lysosomal toxicities and biogenesis of TNTs, mitochondrial toxicities were observed through morphological changes in the early time points (3h and 6h) using MitoProbe JC-1 dye which measures mitochondrial membrane potential (ΔΨ_M_). Lipophilic, cationic JC-1 dye can enter healthy mitochondria and aggregate (emission @590 nm in the red channel) inside, whereas, in unhealthy mitochondria with decreased membra.ne potential, the dye crosses in and out through the relatively open membrane pores and stays in monomers (emission @527 nm in the green channel). Adding to our previous observations, we noticed a decrease of ΔΨ_M_ transiently in the early time points (3h and 6h) in primary astrocytes, at the later time point (12h and 24h) mitochondria are healthier and rescued from protofibrils induced toxicities (Fig 4A and B). The quantification shows JC-1 labelled red coloured healthier, larger mitochondria in the control and recovered healthier cells after 24h (Fig 4C). The toxic mitochondria in early time points (3h and 6h) were observed to possess shorter branch length, compared to the control and the recovered cells at later times (12h and 24h) (Fig 4D). Mitochondrial toxicities were also observed in U-87 MG cells through morphological changes using mitotracker staining (Fig 4E, upper panel) and mitochondrial membrane potential (ΔΨ_M_) using MitoProbe JC-1 (Fig 4E, lower panel). Similar to our results in primary astrocytes, images of U-87 MG cells show that at the early time points (3h and 6h) mitochondria are smaller sized, fragmented in morphology (Fig 4E, upper panel) and quantifications show increased JC-1 stained greener mitochondria or mitochondria with decreased ΔΨ_M_ (Fig 4F). Thus, it is evident that primary astrocytes and U-87 MG cells adapt and are rescued from α-SYN protofibril-induced transient mitochondrial toxicities at later times.

**Figure 4:**
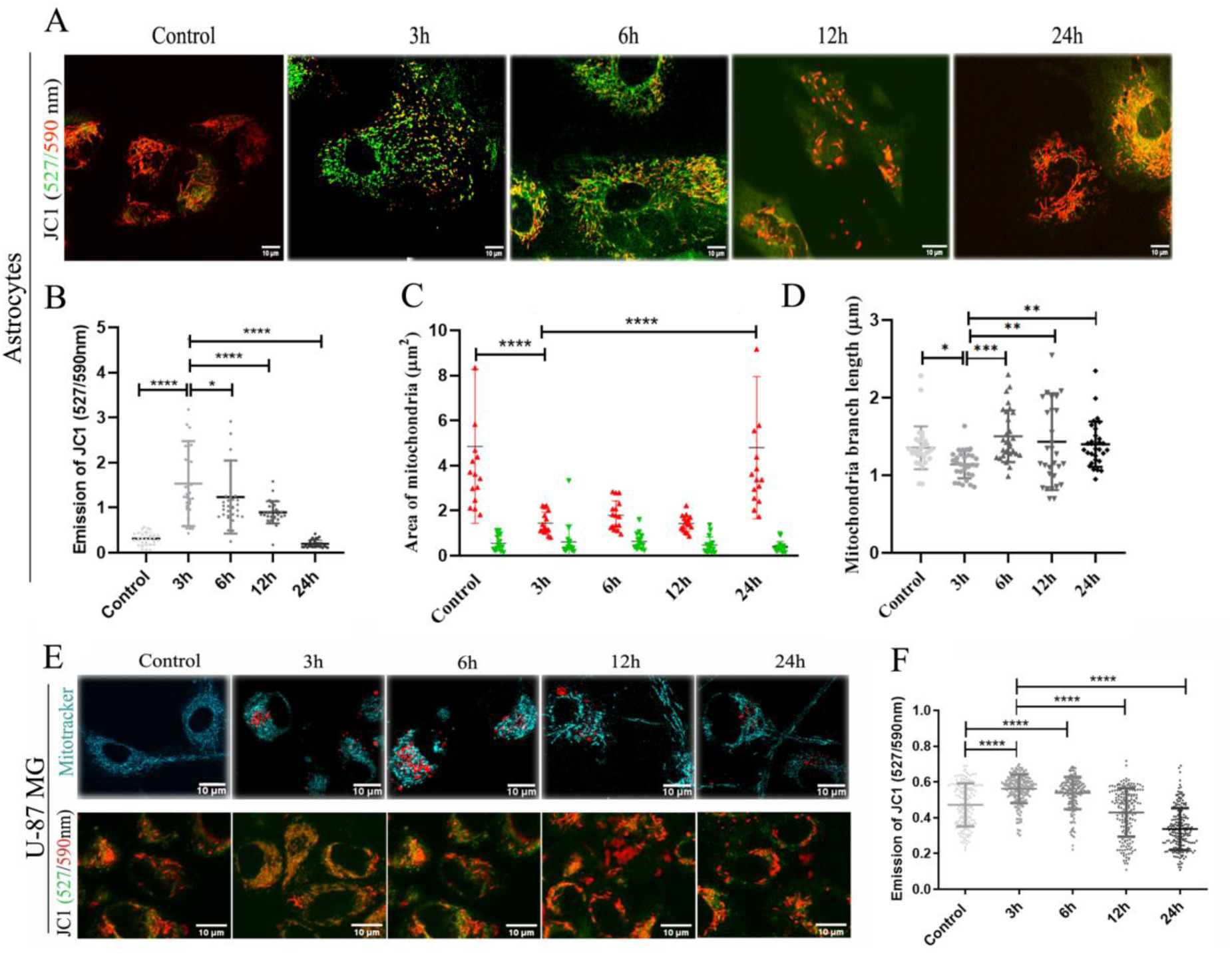
Effect of α-SYN protofibrils on mitochondria in astrocytes and U-87 MG cells. Primary astrocytes and U-87 MG were treated with 1µM α-SYN protofibrils for 3h, 6h, 12h and 24h and mitochondrial membrane potentials were detected using JC-1 staining (emission ratio of 527/590 nm). A) Primary astrocytes stained with JC-1. B) Quantification of green vs red intensities, C) Size of green/red mitochondria and D) mitochondria branch length by MiNA analysis were quantified from JC-1 astrocyte images. E) U87MG cells treated with 1µM α-SYN protofibrils for 3-24 h, were stained with mitotracker (top panel -cyan) and JC-1 (bottom panel) to visualize mitochondrial morphology. F) Quantification of green vs red intensities from JC-1 U-87 MG images. Quantification of green vs red intensities. Quantifications are done from 10-15 image frames of a set and analysing 10-20 cells from each image. Scale bars are denoted on the images. Data are expressed as mean ± SD, *** p ≤ 0.001. Statistics were analysed using two-way ANOVA. n=3.

### Cell-to-cell transfer of mitochondria in α-SYN protofibril treated astroglia cells

The transfer of mitochondria between α-SYN protofibrils-treated U-87 MG cells was further examined by co-culturing these cells (Fig 5A). MitoDsRed (red) and EGFP-lifeact (green) transfected U-87 MG cells were co-cultured and treated with α-SYN protofibrils for 3h, 6h, 12h and 24h. Results show mitoDsRed labelled red mitochondria in the green EGFP-lifeact stained cells upon treatment with α-SYN protofibrils (Fig 5A). The transferred red mitochondria in the green cells were observed maximum at early time points (3h and 6h), when cells show enhanced mitochondrial toxicities and increased numbers of TNTs (Fig 5B). When conditional media from the mitoDsRed transfected cells were given to the green cells transfected with EGFP-lifeact, we did not observe any significant transfer of mitochondria. The result excludes the probability of transfer of mitochondria through exosomes. The morphology of the transferred mitochondria in the green cells mostly appear round in shape rather than elongated and healthy. Quantification shows transferred mitochondria are smaller in size and round shaped, compared to the mitochondria of the control cells (Fig 5C). Therefore, we further studied fate of cell-to-cell transfer of toxic mitochondria, following ROS levels in the cells upon protofibril treatment at different time intervals. We also observed increased total cellular ROS at an early time point (3h) and gradual decrease in ROS levels at later time points (6h, 12h and 24h) in U-87 MG cells (Fig 5D and E). Fluorescence intensity of DCFDA was analysed using flow cytometer to quantify the cellular ROS levels (Fig 5E). The quantification of DCFDA flow cytometer data show similar results in primary astrocytes, significant increase in cellular ROS levels at 3h compared to the control cells, and then decrease at later time points (6h, 12h and 24h) (Fig 5F and G).

**Figure 5:**
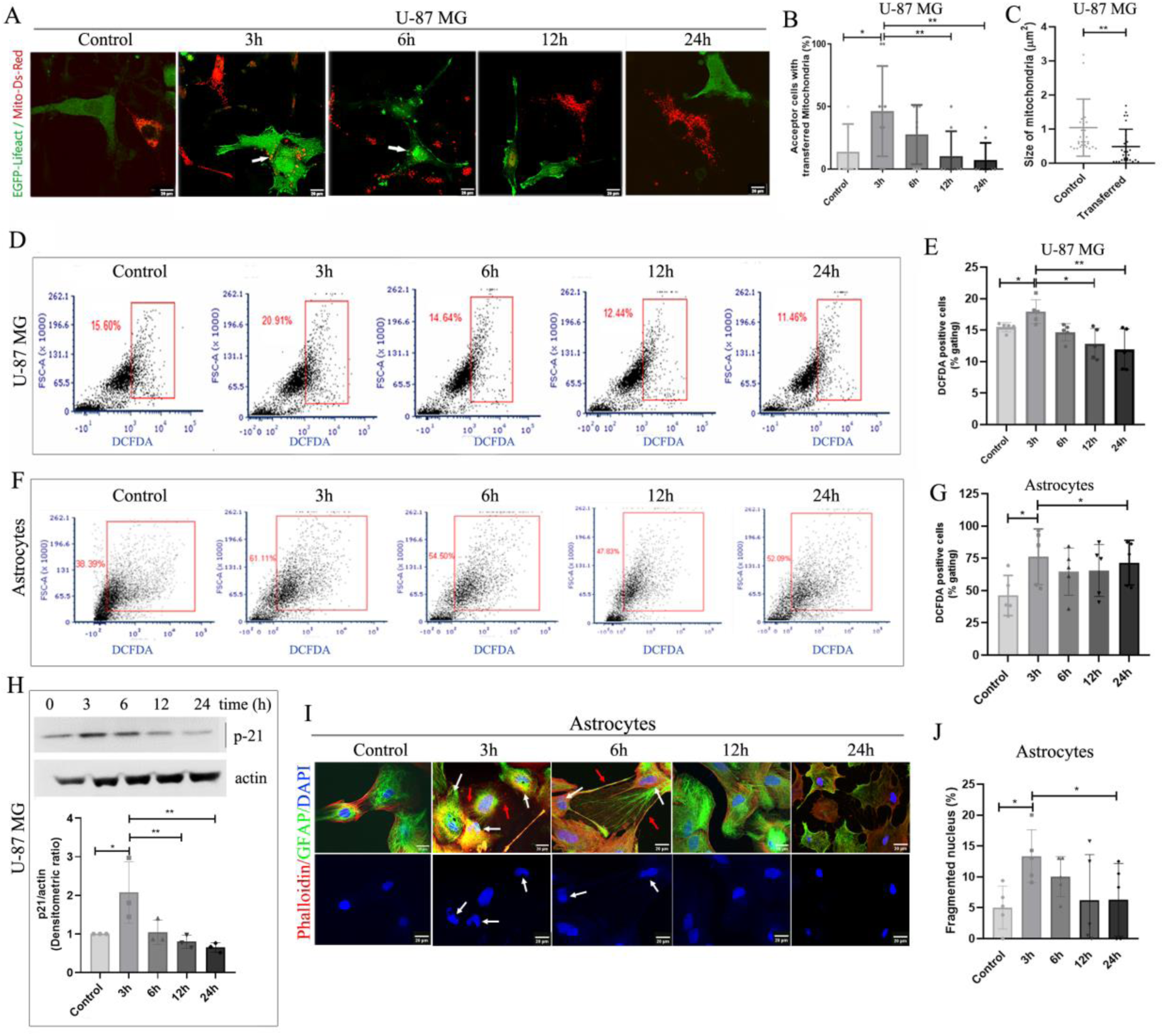
Cell-cell transfer of mitochondria and α-SYN-induced ROS mediated cellular senescence. A) Transfer of mitochondria was detected in co-culture experiment. U87MG cells were co-cultured using transiently transfected with Mito-ds-Red (red) and EGFP-lifeact (green) respectively and treated with 1µM α-SYN protofibrils for 3h-24h. Confocal microscopy images show the transfer of Mito-ds-Red labelled mitochondria to EGFP-lifeact labelled green cell population. B) Quantification of percentage of acceptor cells with transferred mitochondria. C) quantification of size of the transferred mitochondria vs the control mitochondria. D, F) Estimation of total cellular ROS by DCFDA assay in the cells treated with 1µM α-SYN protofibrils for 3h-24h by flow cytometry in U-87 MG and astrocytes respectively. E, G) Quantification of median fluorescence intensity obtained from the same. Quantifications are done from 5 image frames of a set and each image frame have 10-15 cells. H) Western blot analysis to estimate the levels of senescence marker p21 in the cells treated with 1µM α-SYN protofibrils. I) In astrocytes images white arrows indicate DAPI stained fragmented nucleus while the red arrows indicate formation of TNTs from those cells. J) Quantification of percentage of fragmented nucleus on α-SYN protofibrils treatment. Quantifications are done from around 10 image frames of a set and each image frame have 10-20 cells. Scale bars are denoted on the images. Data are expressed as mean ± SD, *** p ≤ 0.001. Statistics were analysed using two-way ANOVA except graph C which was analyzed by student t-test. n=3.

### α-SYN protofibril-induced ROS mediated cellular senescence and its correlation with transient biogenesis of TNTs

Increased levels of cellular ROS play a significant role to induce cellular senescence and reduction of ROS accumulation can reverse p-21 mediated cellular senescence (Macip, Igarashi et al. 2002). Studies have also shown critical role of α-SYN induced toxicities in cellular senescence associated neurodegeneration (Verma, Seo et al. 2021, Miller, Campbell et al. 2022). α-SYN toxicities induced-increased ROS levels and its elimination in correlation to TNT biogenesis, motivated us to look into the senescence states in astroglia cells (Macip, Igarashi et al. 2002). We have observed transient increase in p-21 levels with α-SYN protofibrils treatment in western blot analysis at earlier time point (3h) in U-87 MG, then p-21 levels decrease gradually over time (6h, 12h and 24h) (Fig 5H). Further, we have observed irregular nucleus, invaginations and fragmented nucleus during the transient time window 3h and 6h, which corresponds to the α-SYN protofibrils induced transient toxicities and biogenesis of TNTs in primary astrocytes (Fig 5I-J) and U251 cells (Fig EV3A and B). We established cells undergo transient cellular senescence at early time point (3h) by measuring β-galactosidase activity as the senescence marker (Fig EV3C). It is known that DNA damage related nuclear size irregularities often associated with hyperproduction of cellular ROS and mediated senescence. The transient cellular senescence corresponds to transient biogenesis of TNTs at early hours in α-SYN treated cells. Eventually, reduction of ROS accumulation caused reversal of p-21 dependent cellular senescence at later times.

### Clearance of α-SYN induced toxicities in cell survival and proliferation

It is a well-established fact that extracellularly applied α-SYN protofibrils are toxic to neurons and cause gradual neuronal death (Li, Yuan et al. 2021). Conversely, we observed astrocytes and astrocytes origin cancer cells survive by rescuing α-SYN protofibrils induced toxicities. We also observed that biogenesis of TNTs and cell-to-cell transfer precedes clearance of α-SYN protofibrils induced toxicities. Recent studies have shown that, cell-to-cell transfer of mitochondria in astrocytes *via* TNTs facilitate cell survival (Valdebenito, Malik et al. 2021). Therefore, we investigated cell viability using α-SYN protofibrils in primary astrocytes, U-87 MG, and U251 astroglia cells. MTT assay measures cell metabolic activity and is used as an indicator for cell viability. The results pertaining to this assay shows concentration dependent increase of cell viability upon treatment of toxic α-SYN protofibrils at later time points (12h and 24h) in astrocytes (Fig 6A), compared to the respective controls and early time points (3h and 6h). α-SYN protofibrils induced toxicities at earlier times also resulted in decreased metabolic activity (Fig 6A). α-SYN protofibrils treated U-87 MG (Fig 6B) and U251 (Fig 6D) cells show proliferations at later times (12h and 24h) compared to their respective controls. U-87 MG cells also show concentration dependent proliferations at later times (24h) (Fig 6C). Further, we manually counted the cells to reconfirm that the increasing concentrations of toxic protofibrils caused increase in cell numbers or cell proliferations, after 24 and 48 h of treatments in astroglia cells (Fig EV4A and B). Overall, the results showed that, instead of developing progressive toxicities or cell death overtime with α-SYN protofibrils treatment, the astroglia cells adapt to overcome the stress and proliferate during post recovery time. We have verified the toxicity of the protofibrils in the neurons derived from differentiation of neuroblastoma (N2a) cells. The results show time- and concentration-dependent toxicities and cell death in the treated cells (Fig EV4C and D).

**Figure 6:**
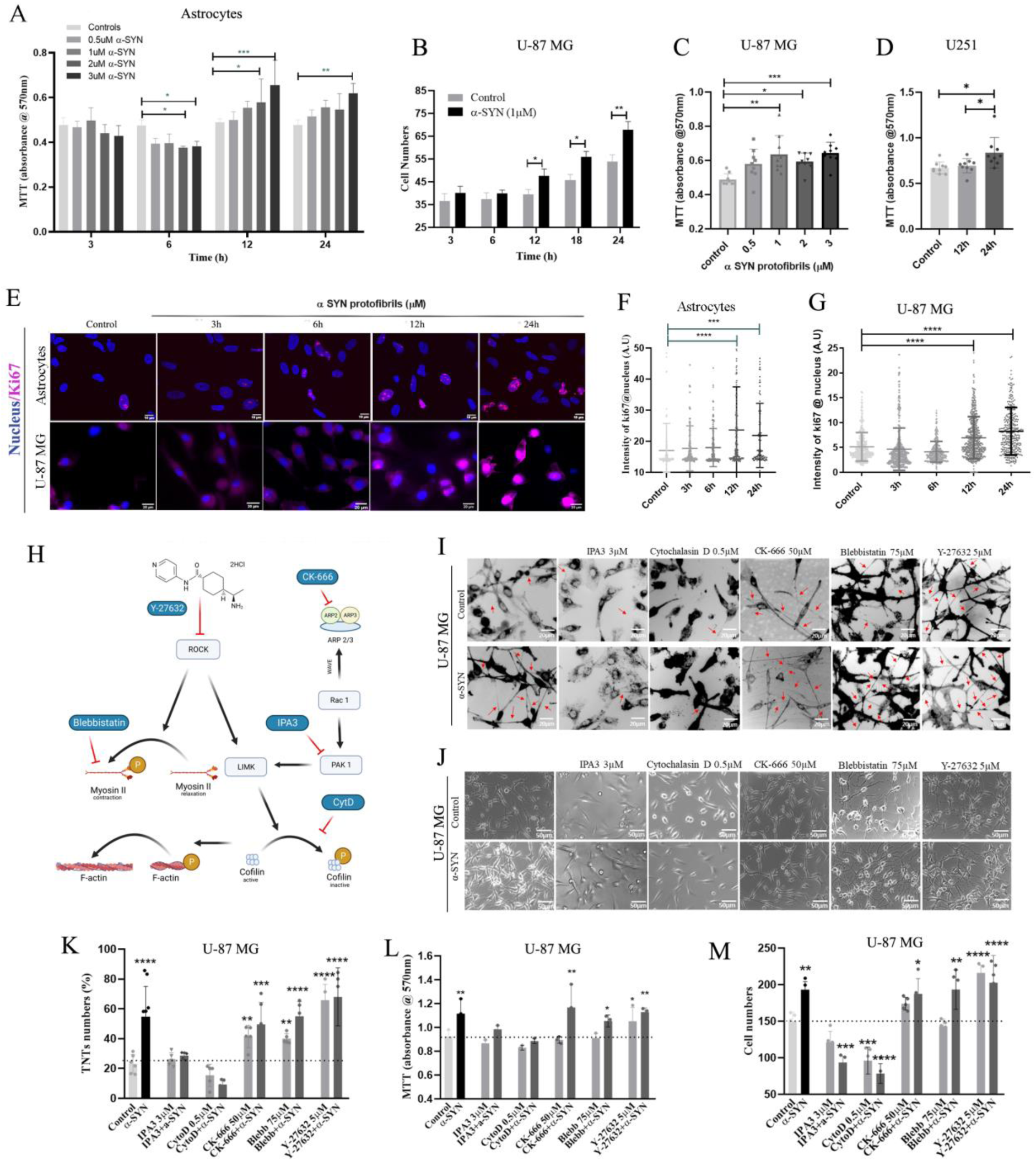
TNT biogenesis pathways in cell proliferation. A) Absorbance measured at 570 nm after MTT assay performed in astrocytes treated with varying concentrations (0.5µM, 1µM, 2µM and 3µM) of α-SYN protofibrils at 3h, 6h, 12h and 24h. B) Cell number quantification of U-87 MG cells treated with 1µM α-SYN protofibrils after 3h, 6h, 12h, 18h and 24h. C) MTT absorbance estimated at 24h on treatment with varying concentrations (0.5µM, 1µM, 2µM and 3µM) of α-SYN protofibrils in U-87 MG cells. D) Estimation of MTT absorbance at 12h and 24h after treatment with 1µM α-SYN protofibrils in U251 cells. E) Fluorescence images of astrocytes and U87MG cells treated with 1µM α-SYN protofibrils for 3h-24h were stained with proliferation marker Ki67 along with nuclear stain. F, G) Quantification of the intensity of Ki67 per cell in astrocytes and U-87 MG respectively. Quantifications are done from 15 image frames of a set and each image frame have 20-25 cells. H) Flow chart depicting mode of action of actin inhibitors. I) DiD (membrane dye) stained, control and 1µM α-SYN protofibrils treated U-87 MG cells pre-treated (before 30min) with 3 µM IPA3, 0.5 µM cytochalasin D, 50 µM CK-666 (Arp2/3 inhibitor), 75 µM Blebbistatin and 5 µM Y-276325 (ROCK inhibitor). Red arrows indicate formation of TNT-like structures and K) Shows quantification of TNT numbers. J) representative images showing cell numbers with the above-mentioned treatment and L, M) quantification of MTT absorbance and cell numbers of the same respectively. Quantifications are done from 5 image frames of a set and each image frame have 15-20 cells. Scale bars are denoted on the images. Data are expressed as mean ± SD, *** p ≤ 0.001. Statistics were analysed using two-way ANOVA, only the graph 6B was analysed using one-way ANOVA. n=3.

Since we have observed significant increase in cell numbers and cell viability upon α-SYN protofibrils treatment, we checked for cell proliferation using Ki67 as the marker. Astrocytes and U-87 MG cells immunostained with Ki67 antibody and nuclear stain DAPI (Fig 6E) show a significant increase in Ki67 overexpression in the nucleus at 12h and 24h in comparison to control and early time points (3h and 6h) (Fig 6F and G). Overall, the results indicate toxic α-SYN induced initial stress (till 6 h) promotes biogenesis of TNTs and cell-to-cell transfer of organelles, consequences of which aid to rescue the cells from organelle toxicities or cellular stresses. Further, to deal with recovery, cells probably facilitate proliferation.

### TNTs biogenesis pathways in astroglia proliferation

TNTs are structurally open-ended membrane actin conduits. Thereby, it is obvious that modulation of membrane and cytoskeleton will play a major role in their biogenesis. However, the exact mechanism of TNT biogenesis is not known. Studies on screening of inhibitors in the actin signalling pathways, could unfold molecular events behind the α-SYN toxicity induced biogenesis of TNTs. Therefore, we studied different actin inhibitors (Fig 6H) to see their effects on biogenesis of TNTs at early time (3h) (Fig 6I) and cell proliferation at later time (24h) in U-87 MG cells (Fig 6J). We observed that Cytochalasin D (inhibits actin polymerization and interaction of G-actin-cofilin) and IPA-3 (PAK1/2 inhibitor) inhibited α-SYN protofibril induced TNTs formation, whereas, CK-666 (Arp2/3 inhibitor), Blebbistatin (myosin-II-specific ATPase inhibitor), and Y-27632 (ROCK inhibitor) promote biogenesis of TNTs (Fig 6K). Similar to the reports of earlier studies (Henderson, Ljubojevic et al. 2022), we have also observed that Arp2/3 inhibitor CK-666 caused formation of significantly longer TNTs (Fig 6I and EV5).

To understand the role of TNTs in cell proliferation, cell viability was assayed using MTT (Fig 6L) and cell numbers were determined by manual counting (Fig 6M). TNT-numbers (Fig 6K) and cell proliferation (Fig 6L and M) data showed that the actin inhibitors, Cytochalasin D, and IPA-3, which prevent biogenesis of TNTs, inhibit α-SYN protofibrils induced cell proliferation as well (Fig 6L and M). The actin inhibitors, CK-666 and Blebbistatin which facilitate biogenesis of TNTs, did not alter α-SYN protofibrils induce.d proliferation. However, the inhibitors alone (CK-666, and Blebbistatin) did not show significant effect on cell proliferation. We observed the ROCK inhibitor Y-27632 which works as an intriguing factor in biogenesis of TNTs, both independently and in presence of α-SYN protofibrils significantly increased proliferation (Fig 6L and M).

### Biogenesis of α-SYN induced TNTs through modulation of ROCK pathway, resulting in increased cell survival and proliferation

Recent study has shown that, the ROCK inhibitor (Y-27632) is an intriguing compound that boosts biogenesis of TNTs *via* Myosin II mediated actin remodulation (Scheiblich, Dansokho et al. 2021). However, it is not yet known how α-SYN dominates modulation of ROCK inhibition signalling mediated TNT biogenesis, over the cofilin-G-actin interaction (Cytochalasin-D inhibited pathway) mediated actin remodulation. To unfold the mechanism, we followed internalization of α-SYN-TMR protofibrils in presence of Cytochalasin-D and ROCK inhibitor (Y-27632) using flowcytometry quantification (Fig 7A). We observed Cytochalasin-D inhibited internalization of α-SYN protofibrils, however, Y-27632 did not show any effect (Fig 7B).

**Figure 7:**
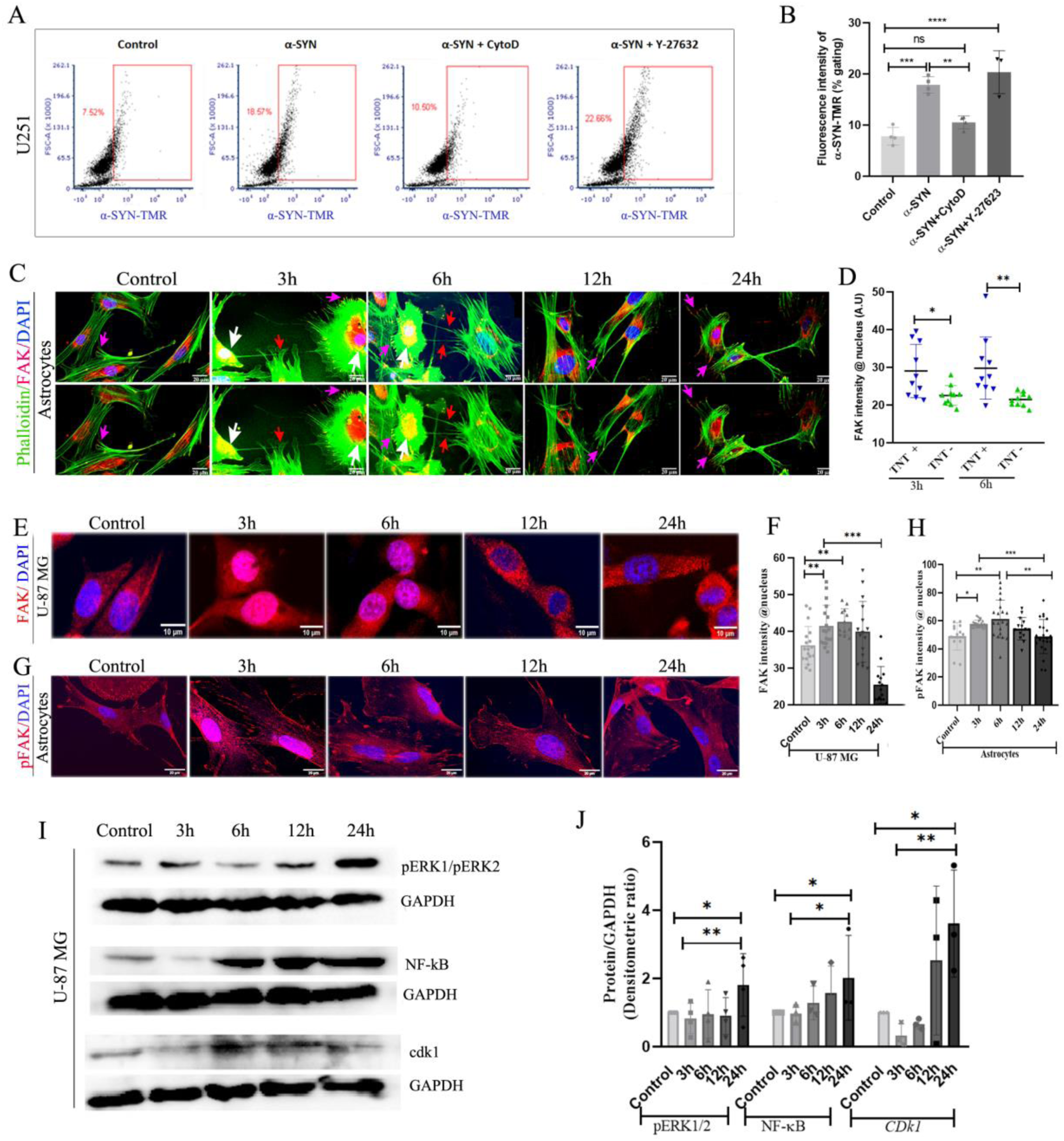
FAK translocation, ROCK remodulation and activation of proliferation pathway. A) Estimation of uptake of α-SYN-TMR protofibrils by U251 cells treated with 1µM α-SYN protofibrils and pre-treated (before 30min) with 0.5 µM cytochalasin D and 5 µM Y-276325 (ROCK inhibitor) by flow cytometry. B) Quantification of percentage gated fluorescence intensity of α-SYN-TMR protofibrils in the above experiment. C) Fluorescence images of maximum intensity projected confocal z-stacks in astrocytes stained with Phalloidin (green), FAK (red). Upper panel with nucleus stained with DAPI and lower panel without DAPI. White arrows indicate nuclear colocalization of FAK, red arrows indicate TNTs and pink arrows indicate FAK at focal adhesion. D) Quantification of FAK intensity in the nucleus of the cells with and without TNTs. E, G) Fluorescence images of maximum intensity projected confocal z-stacks in U-87 MG cells stained with FAK (red) and pFAK (red) with DAPI respectively. F, H) Quantification of FAK and pFAK fluorescence intensity per cell in the nucleus. Quantifications are done from 15 image frames of a set and each image frame have 15-20 cells. I) Western blot images showing increased intensity of pERK1/pERK2, NF-κB and cdk1 with time on 1µM α-SYN protofibrils treatment in U-87 MG cells. J) Quantification of western blots. Scale bars are denoted on the images. Data are expressed as mean ± SD, *** p ≤ 0.001. Statistics were analysed using two-way ANOVA. n=3.

### α-SYN-induced cellular senescence results in translocation of FAK in the nucleus

Then, we tried to understand how α-SYN protofibrils inhibits ROCK signalling pathway to promote TNTs biogenesis and cell proliferation. Our results showed α-SYN protofibrils induce nuclear deformity and cellular senescence in astroglia cells at early time points (3h and 6h) (Fig 5H-J). This is evident from several studies that, inhibition of focal adhesion kinase (FAK) induces the DNA-damage and nucleus deformity that accompanies cellular senescence (Chuang, Wang et al. 2019, Zhou, Yi et al. 2019). Loss of integrin-dependent cell adhesion in cellular stress modulates FAK and facilitates its translocation from plasma membrane to enter to nucleus (Lietha, Cai et al. 2007, Lim, Chen et al. 2008, Lim, Miller et al. 2012). We observed de-adhesion of astroglia cells (U-87 MG) with α-SYN protofibrils treatments at early time points (3h and 6h) when cells show toxicities and TNTs biogenesis, compared to the control and cells at later times (12h and 24h) (Supplementary Fig EV6A). De-adhesion of cells displaces FAK localization from the focal adhesion sites, which leads to modulation of ROCK signalling cascade (Pirone, Liu et al. 2006). Therefore, we followed FAK translocation with α-SYN protofibrils treatments and its internalization. We observed treatments with α-SYN protofibrils cause nuclear translocation of FAK in the astrocytes that are connected by TNTs at early time points (3h and 6h) (Fig 7C and D). Presence of nuclear FAK was verified by 3D images at xz and yz planes (Fig EV6B). We also observed treatments with α-SYN protofibrils cause nuclear translocation of FAK to the nucleus in the U-87 MG (Fig 7E and F) and U251 (Fig EV6C and D) cells at early timepoints (3h and 6h), whereas, at later timepoints with recovery from α-SYN toxicities FAK relocates back to PM and cytosol. We have also observed nuclear translocation of activated phospho-FAK (Tyr 397) transiently at early time points (3h and 6h) in astrocytes (Fig 7G and H).

### Nuclear translocation of FAK modulates ROCK pathway to induce TNT biogenesis and cell proliferation

Inhibition of integrin-mediated FAK activation prevents ROCK activation, and inhibits cell adhesion. Moreover, when active/inactive FAK is displaced from cell-adhesion sites in non-adherent cells, then Rho mediated activation of ROCK kinase maintains cytoskeleton tension by regulating actin re-modulation (Pirone, Liu et al. 2006). Probably, FAK translocation dynamics in α-SYN treated toxic cells is driving ROCK inactivation in early timepoints, and subsequently activation at later timepoints to restore cytoskeleton tension. Actin re-modulation in rescued cells, may result in increased cell proliferation. In the western blots we have determined increased levels of ROCK mediated cell proliferation markers ERK1/2, NF-κB and cdk1 (Fig 7I and J). Overall, the results delineate α-SYN protofibrils induced toxicities may modulate ROCK inhibitory pathway to promote biogenesis of TNTs in the astroglia cells for a transient time at early hours after the treatment. The rescued cells post α-SYN treatment and transient TNT biogenesis, eventually may re-activate ROCK signalling to restore cytoskeleton tension, and resulting in enhanced cell proliferation.

## Discussion

PD pathology develops with the toxic burden of α-SYN aggregates in the brain (Kalia and Lang 2015). Even though cytoplasmic inclusion of α-SYN in the neurons is the central hallmark of neuropathology development in PD. Evidence suggests release of α-SYN aggregates from degenerated neurons in the extracellular brain play a significant role in PD pathology. Role of extracellular α-SYN in cell-to-cell transfer and PD pathology progression have widely been studied in several model systems (Neupane, De Cecco et al. 2022). Recent studies have shown that extracellular aggregates, taken up by astrocytes and microglia, facilitate clearance of aggregates by promoting TNTs-mediated glial cross-talks and cell-to-cell spreading (Rostami, Holmqvist et al. 2017, Scheiblich, Dansokho et al. 2021). Astrogliosis and microgliosis act as essential mediators in maintaining cellular homeostasis in the PD brain (MacMahon Copas, McComish et al. 2021). On the other hand, studies have shown accumulations of α-SYN in glial population in several synucleinopathies (Mavroeidi and Xilouri 2021). Expressions of α-SYN have also been implicated in astrocytoma-glioblastoma brain tumours, by immunohistological staining of the patient tissues (Bruening, Giasson et al. 2000, Fung, Rorke et al. 2003).

The contribution of neurons and glial cross-talks and their dynamic interplay in clearance of neurodegenerative aggregates to prevent pathology development remain unexplained. In this context, we observed transient biogenesis of TNTs in astrocytes and astroglia cells (U-87 MG and U251) in response to toxic burdens of α-SYN, which corresponds to transitory toxicity levels of lysosomes and mitochondria. We have also observed a significant but transient increase in cellular ROS, which corresponds to transient biogenesis of TNTs. The transient biogenesis of TNTs facilitates cell-to-cell transfer of lysosomes accumulated with toxic α-SYN protofibrils and toxic mitochondria. Several studies in various models have also shown that oxidative stress and ROS promote biogenesis of TNTs (Raghavan, Rao et al. 2021). A recent study has shown an on-demand formation of TNTs to clear toxic α-SYN protofibrils and alleviate ROS levels in microglia (Scheiblich, Dansokho et al. 2021).

Increased ROS levels directly affects mitochondrial dysfunction, DNA damage, and increased protein expressions related to ageing (Davalli, Mitic et al. 2016). Mitochondrial dysfunction and DNA damage play key roles to induce cellular senescence (Gallage and Gil 2016). Evidence showed α-SYN protofibrils induce senescence in reactive astrocytes (Verma, Seo et al. 2021). In this study, we have observed α-SYN protofibrils induced senescence in astroglia cells and the senescent astrocytes possess characteristics of fragmented nucleus. We have observed α-SYN protofibrils induced mitochondrial abnormality, fragmented irregular nucleus, increased β-galactosidase activity, and p21-pathway dependent premature cellular senescence for a transitory time period. The astroglia cells recover from senescence related protein expressions, nuclear fragmentation and morphological changes, after transient biogenesis of TNTs and TNT-mediated cell-to-cell transfer. ROS induced DNA damage and mitochondrial toxicity do not always lead to apoptosis (Borges, Linden et al. 2008). Cells activate repair mechanism/s in case of delayed apoptosis. Similarly, reduction of ROS accumulation can reverse p21-mediated stress induced premature senescence (Macip, Igarashi et al. 2002). Thus, α-SYN protofibrils induced transitory increase of ROS levels and subsequently its reduction in correlation to transient biogenesis of TNTs, could explain the reversing of oxidative stress-induced premature cellular senescence in astroglia cells.

The first report of mitochondrial transfer through TNTs was in 2006 (Spees, Olson et al. 2006). Later, formation of TNTs and transfer of mitochondria through them were reported by many studies in several pathophysiology conditions, where transfer of healthy mitochondria resulting in rescue of cells from apoptosis or toxicities (Lou, Fujisawa et al. 2012, Han and Wang 2021). Studies have shown that transfer of healthy mitochondria from mesenchymal stem cells (MSCs) improves health of damaged acceptor cells (Ahmad, Mukherjee et al. 2014, Tseng, Lambie et al. 2021). However, it is not the case that TNTs transfer only healthy mitochondria to neighbouring cells. Microglia and astrocytes are reported to facilitate dilution of its toxic burdens by transferring toxic α-SYN aggregates and mitochondria by sharing with its neighbours through TNTs (Loria, Vargas et al. 2017, Scheiblich, Dansokho et al. 2021). Another recent study demonstrated that neuronal cells transfer toxic mitochondria to microglia via TNTs, whereas microglia protects neurodegeneration by transferring healthier mitochondria to neurons (Chakraborty, Nonaka et al. 2023). We observed α-SYN protofibrils induced increased ROS production at early time points (3h and 6h), caused dysfunctional mitochondria with reduced mitochondrial membrane potential, resulting in generation of fragmented fission mitochondria and disintegration of mitochondrial network. In the co-culture experiment we have observed that cells transfer toxic fragmented mitochondria to its neighbouring cells at early hours after α-SYN protofibrils treatment. In our astroglia cellular models, we observed only unidirectional transfer of organelles (α-SYN protofibrils accumulated lysosomes, lysosomes and mitochondria) at early time points, when cells showed visible lysosomal and mitochondrial toxicities. The observations suggest that astrocytes and astroglia cells share their toxic organelles with surrounding neighbours to dilute the toxic burdens, probably to continue survival under pathological stress. TNTs are predominantly observed in the cell types which possess inherent anti-apoptotic properties, like primary neurons, neuronal cells, neuroglial cells and cancer cells (Rustom, Saffrich et al. 2004, Gousset, Schiff et al. 2009). Probably, TNTs act as an alternative pathway to get rid of its toxic burdens to facilitate degradation by diluting their toxic content through spreading, so the cells can sustain the survival condition under pathological stress.

Cell-to-cell transfer of neurodegenerative aggregates through TNTs may not be a cellular survival mechanism for neurons, even though, neurons possess inherent anti-apoptotic properties like other brain cells (Tardivel, Begard et al. 2016, Sharma and Subramaniam 2019). We observed α-SYN protofibrils treatments eventually lead to cell death in mitotically incompetent neurons, whereas astrocytes and astroglia cells enhance proliferation and cell survival. Cell-to-cell transfer of neurodegenerative aggregates drives spreading of aggregates, which acts as seeds for further propagation in neurons, whereas, neuroglial cells and cancer cells rescue cellular toxicities (Nath, Agholme et al. 2012, Rostami, Holmqvist et al. 2017). Recent study in microglia has shown, rescue of toxic burdens of α-SYN *via* TNT-mediated transfer of toxic aggregates to its healthier neighbours and borrowing of healthier mitochondria (Scheiblich, Dansokho et al. 2021). We have observed in our astroglia models that α-SYN treatment promotes biogenesis of TNTs and cell-to-cell transfer of toxic organelles preceding the clearance of toxic burdens. However, the study cannot completely rule out the role of inherent conventional clearance mechanisms in the rescue of toxic burdens in astrocytes. Otherwise, biogenesis of TNTs could be an alternative aid to maintain cellular homeostasis during rapid clearance process in response to massive toxic burdens. Nevertheless, we have established strong correlation between transient biogenesis of TNTs in relation to α-SYN-induced toxic burdens.

Therefore, the important question that needs to be understood is how biogenesis of TNTs and clearance of cellular toxicities associate to proliferation? Cell proliferation following transient TNT biogenesis in response to massive toxic burdens could also be the consequence of maintaining cellular homeostasis and cell survival. Several studies have indicated that biogenesis of TNTs may induce cell proliferation, to protect the cells from cell death under pathogenic conditions and chemo- or radio-therapy related stress (Osswald, Jung et al. 2015, Han and Wang 2021, Wang, Chen et al. 2021, Saha, Dash et al. 2022). The studies show mostly limited or indirect correlation between TNTs and cell proliferation. We have observed that α-SYN protofibrils treated cells eventually trigger cell proliferation in astroglia cells, during post recovery time.

Our results on small molecule actin modulators indicate ROCK inhibitory pathway mediated biogenesis mechanism/s of TNTs and its correlation with enhanced proliferation. Small molecules actin modulators (Cytochalasin D, and IPA-3) that inhibit biogenesis of TNTs, prevent α-SYN protofibril induced cell proliferation. On the other hand, the actin modulators CK-666 and Blebbistatin, which facilitate biogenesis of TNTs, did not show significant effect on the enhancement of cell proliferation. Even though these actin inhibitors (CK-666 and Blebbistatin) show better cell survival compared to the molecules that inhibit TNTs. Only ROCK inhibitor Y-27632 mediated biogenesis of TNTs significantly promotes proliferation, similar to proliferation induced by α-SYN protofibrils. ROCK inhibitor Y-27632 inhibits LIMK dependent cofilin-G-actin interaction and inhibits actin polymerization. Whereas, IPA-3 and Cytochalasin-D could also inhibit actin polymerization through LIMK pathway (Scheiblich, Dansokho et al. 2021). However, we observed ROCK inhibition promotes biogenesis of TNTs, whereas, IPA-3 and Cytochalasin-D inhibit TNTs. Our result suggests, ROCK inhibition may modulate TNTs biogenesis by regulating actin cytoskeleton through myosin-II-specific downstream signalling molecules, which dominates over cofilin-mediated actin modulation. Similarly, a recent study (Scheiblich, Dansokho et al. 2021) has shown that α-SYN protofibrils promote biogenesis of TNTs in microglia cells *via* regulating ROCK inhibitory pathway by phosphorylation of myosin light chain phosphatase (MLCP).

Thus, we tried to unfold how α-SYN protofibrils induced actin-remodulation may modulate ROCK inhibitory signalling cascades to induce TNT biogenesis. We have found that α-SYN induced transient senescence related toxicities caused nuclear localization of FAK and pFAK for transitory period, which corresponds to TNT biogenesis at early timepoints (3h and 6h). Organization of FAK at focal adhesion sites of PM regulates activation of Rho mediated ROCK signalling, and its de-localization from focal adhesion inhibits ROCK pathway (Schober, Raghavan et al. 2007). On the other hand, in low adhesion conditions displacement of inactive/active FAK/pFAK from focal adhesion sites leads to Rho kinase mediated activation of ROCK signalling, to maintain cytoskeletal tension (Pirone, Liu et al. 2006). Then, restoring of cytoskeleton tension may re-translocate FAK. ROCK activation enhances cell proliferation *via* ERK1/2 and NFκB activation. In this line, we have detected α-SYN toxicities induced FAK translocation dynamics, which could modulate transient inhibition of ROCK signalling to induce biogenesis of TNTs at early times and later may activate to restore cytoskeleton tension. We have also seen proliferation of astroglia cells *via* activation of ERK1/2 and NFκB signalling during TNT-mediated post recovery time (12h and 24h). Thus, our results suggest that biogenesis of TNTs is not the root cause of cell proliferation, rather modulation of α-SYN protofibrils induced ROCK mediated actin-regulatory signalling pathway related to biogenesis of TNTs triggers cell proliferation.

α-SYN inclusions in the astroglia cells and glial inclusion or fibrous gliosis in the areas of neurodegeneration are widespread in PD (MacMahon Copas, McComish et al. 2021). Moreover, extracellular release of α-SYN promotes proliferation of glioma cells (Gibson, Purger et al. 2014, Venkatesh,Johung et al. 2015, Venkatesh, Morishita et al. 2019). Thus, the study is important for understanding the contributions of α-SYN toxicities in glial cells proliferation. Our results further reveal relevance of glial crosstalk in clearance of neurodegenerative proteins, and its implication in astroglia cells proliferation. The astrocytes and microglia drive clearance of toxic aggregates via TNT-facilitated cell-to-cell transfer, which play significant role in the modulation of neuroinflammation and pathology progression (Rostami, Holmqvist et al. 2017, Rostami, Mothes et al. 2021, Scheiblich, Dansokho et al. 2021)

In conclusion, our study reveals α-SYN protofibrils induced biogenesis of TNTs aid to enhance clearance of toxic burdens as a cellular survival strategy to rescue the astroglia cells from ROS induced cell death/cellular senescence. α-SYN protofibrils regulate FAK mediated modulation of ROCK signalling cascades to rescue the cells from toxic burdens by promoting TNT biogenesis. The rescued cells, eventually re-activate ROCK signalling probably to restore cytoskeleton tension and enhance cell proliferation via ERK1/2 and NFκB signalling. TNT-mediated glial cross-talk as an alternative aid to facilitate the cellular clearance under pathological stress will open up new strategy to design therapeutics in PD and other neurodegenerative diseases.

## Materials and methods

### Cell culture maintenance

U-87 MG and U251 cell lines (astrocytoma-glioblastoma origin cancer cell lines) was kind gift from Prof. Kumaravel Somasundaram of Indian Institute of Science, Bangalore, India. Cells were cultured and maintained in DMEM (Gibco #2120395) media supplemented with 10% FBS (fetal bovine serum; Gibco #1600004, US Origin), along with 1% PSN (Penicillin-Streptomycin-Neomycin Mixture; Thermo fisher Scientific #15640055) incubated at 37°C, 5% CO_2._

Neuro 2a (N2a) neuroblastoma cell line was procured from NCCS, India. The cells were cultured and maintained in DMEM media with 10% FBS and 1X Glutamax (Gibco #35050). N2a cells were differentiated with 10 µM Retinoic acid (RA; Sigma-Aldrich #R2625) in the presence of 2% FBS supplemented media for 2-3 days. Differentiation was established from the neurites like morphology.

### Primary astrocytes culture

Five weeks old mice (C57BL/6) were used for primary astrocyte culture. The mice were sacrificed using CO2, and the cerebral cortices were dissected out from mice brain using Olympus SZ51 stereomicroscope. Pieces of cortical tissue were trypsinized using 0.25% trypsin-EDTA solution (Gibco Canada origin #25200-072) for 5 min. The tissue is washed twice in warm HBSS and transferred to Minimum Essential Media (Gibco 61100-087) with 10% FBS (Gibco #1600004, US Origin) containing HEPES (Sigma Aldrich H0887) and D-glucose (Sigma Aldrich G7021). The tissue was triturated using a fire-polished pipette and counted using Trypan Blue in an automated cell counter. Dissociated cells were plated in MEM with 10% FBS on Corning® 100mm TC-Treated Culture Dish at a density of 10 million cells per dish. The media was replaced with fresh MEM with 10% FBS on the next day and maintained in the same for 1 week. After 1 week, cells were trypsinized and re-plated for experiments.

### Purification and Labelling of α-SYN protein

The human α-SYN wild-type (Addgene ID #36046) and α-SYN (wt)-141C (Addgene ID #108866) with a cysteine residue at C-terminus, constructs were purchased and overexpressed in *E. coli*. Then purification was done from periplasmic fraction as described in (Huang, Ren et al. 2005, Jos, Gogoi et al. 2021). Samples were aliquoted and flash frozen in liquid nitrogen and stored immediately at -80°C.

The purified α-SYN (wt)-141C protein was thawed and pre-incubated with reducing agent TCEP (tris(2-carboxyethyl) phosphine-hydrochloride); Sigma Aldrich-C4706; 20mM) to reduce/dissociate any pre-existing cysteine bonds. Then the protein was labelled adding two-fold molar excess of TetramethylRhodamine-5-maleimide (TMR-maleimide; Sigma Aldrich #94506) in 20 mM HEPES buffer (pH7.4). The protein solution was vortexed and the mixture was incubated for 1h at 20°C. Excess label was removed by using PD-10 columns (Himedia #TKC287) (Nath, Meuvis et al. 2010).

### Preparation and characterisation of α-SYN protofibrils

Protofibrils of α-SYN (labelled and unlabelled) was prepared with unlabelled α-SYN wild-type and labelled α-SYN-141C by incubating with 0.65% 4-Hydroxynonenal (10mg/ml stock) (Sigma-Aldrich #393204-1MG) at 37°C for 7 days with moderate shaking (Sackmann, Sinha et al. 2019).

The α-SYN protofibrils formed at the end of 7 days were then characterized by transmission electron microscopy (TEM). The unlabelled and labelled α-SYN protofibrils were resuspended in 1X PBS, coated on a carbon grid with uranyl acetate and dried in a desiccator. The grids were visualized using FEI Tecnai T20 at 200 KV. The α-SYN protofibrils were then lyophilized and stored at -20°C and resuspended in 1X TBS (Tris buffered saline) buffer before experiments.

The toxicity of the protofibrils were assessed by treating the differentiated neuronal cells seeded at a density of 30,000 cells / well in 24 well plate. Cell viability assay was performed using different concentrations of α-SYN protofibrils (0.5µM, 1µM, 2µM, 3µM) over different time period by MTT assay.

### Treatment conditions of astroglia cells by α-SYN protofibrils

All the experiments were carried out by treatment of primary astrocytes, U-87 MG and U251 cells with 1µM α-SYN protofibrils-unlabelled or TMR-labelled for 3h, 6h, 12h and 24h at 37°C, 5% CO_2_ unless mentioned otherwise. The cells were seeded at the same time for all the treated time points and controls. Once cells were adhered properly and appear healthy after 24h of seeding, then treatments were done by adding (α-SYN 1μM) protofibrils in a reverse order of the time points. For example, 24h time points were treated first and then sequentially 12 h, 6 h and 3 h points. Then, experiments were performed for all the time points and controls at a time. Therefore, the condition of the untreated control cells corresponds to all the treatment points.

### MTT assay and cell numbers counting

MTT (3-(4, 5-dimethylthiazol-2-yl)-2, 5-diphenyl tetrazolium bromide) assay is a colorimetric assay used to measure cellular metabolic activity. Astrocytes, U-87 MG and U251cells were seeded at a density of 4000 cells per well in a 96 well plate and incubated overnight at 37°C, 5% CO_2._ Different concentrations of α-SYN protofibrils (0.5µM, 1µM, 2µM, 3µM) were added to the wells over different time period (3h, 6h, 12h, 24h and 48h) and treated with MTT reagent (Himedia # TC191-1G) for 2h. The insoluble formazan crystals were dissolved by adding dimethyl sulfoxide (DMSO; Himedia #TC185) which was later quantified by measuring absorbance at 570 nm using the PerkinElmer-Multimode plate reader spectrophotometer. Cell numbers were counted from the phase contrast microscopy images or DIC images using multi-point option in Fiji (a Java based program developed at the National Institutes of Health, USA).

### Immunocytochemistry (ICC)

After treatment with above mentioned concentrations and time points adding α-SYN protofibrils, the cells were washed with 1X PBS and fixed with 4% PFA. The cells were then incubated with incubation buffer (1mg/ml Saponin and 5%FBS) for 20 min at RT. Respective primary antibodies (as mentioned below) were added and incubated overnight at 4°C, the cells were then washed 3 times with 1X PBS. All the secondary antibody (1:700 dilution) incubations were carried out in dark for 2 hrs at RT, 1X PBS washes were repeated. The coverslips were mounted on glass slides with ProLong Gold antifade reagent with DAPI, (Invitrogen P36941). Alternately, glass bottom 35mm dishes (Cellvis, D35-14-1.5-N) were used in few experiments.

Primary antibodies Ki67 anti-rabbit (Millipore B9260) as cell proliferation marker; FAK Rabbit (CST 3285T) and phospho-FAK ((Tyr397) pAb; Invitrogen 44-624G) as total and active FAK marker; Cathepsin D anti-rabbit (GeneTex #42368), LAMP-1 anti-mouse (BD Biosciences 611042), LAMP-2 anti-mouse (Invitrogen #MA1-205) as lysosomal markers; βIII-tubulin anti-rabbit (Cloud-Clone #PAE711Hu01), Connexin43 Rabbit Ab (CST 3512S), SNCa rabbit pAb (Cloud-Clone #PAB222Hu01), GFAP rabbit pAb (Cloud-Clone #PAA068Mu01) and Phalloidin conjugated with iFlour555 (Abcam #176756) were used to stain TNTs and TMs. Secondary antibodies alexa-488 goat anti-rabbit (A-11070; Invitrogen), alexa-555 anti-mouse (A-1413312; Invitrogen) and alexa-488 anti-mouse IgG (H+L) (A-11059; Invitrogen) were used respectively following the above-mentioned protocol. All primary antibodies were used at a dilution of 1:300, secondary antibodies at 1:500 and phalloidin at 1:700. ICC stained cells were imaged with confocal microscope (Zeiss LSM880, Carl Zeiss, Germany) or fluorescence microscope (IX73-Olympus).

### Western Blot

U-87 MG cells were seeded at a density of 1 million per well in a 6 well plate and were treated with 1µM α-SYN protofibrils from 3h-24h. Post treatment media was removed and the cells were washed with 1X PBS. RIPA buffer was added and the cells were scrapped and collected in 1.5 ml tubes. The tubes were incubated on ice with intermittent vortexing. Later the suspension was spun at 12,000 rpm for 10 min and the supernatant was collected and stored at -20^0^C. After normalizing the protein concentration, western blot assay was performed with primary antibodies GAPDH Mouse (CAB932Hu22 cloud clone dilution 1:1000), NF-kB p65/RelA Mouse mAb (Abclonal A10609 dilution 1:1000), phospho ERK1-T202 + ERK2-T18 (AP0485 Abclonal dilution 1:1000), CDK1 Rabbit pAb (A0220 Abclonal dilution 1:1000), p21 Waf1/Cip1 (12D1) Rabbit mAb (CST #2947 dilution 1:1000) and secondary antibodies, Goat anti-mouse (H+L) (Invitrogen 32430 dilution 1:1500) and Goat Anti-Rabbit (H+L) (Invitrogen 32460 1:1500). The blot was developed with ECL solution (SuperSignal West Femto Trial kit Invitrogen 34094) and were quantified by densitometry using gel analyzer plugin of Fiji software.

### α-SYN internalization

U-87 MG cells were seeded in a 24 well plate at a density of 0.1million/well, pre-treated with 0.5 µM cytochalasin D and 5 µM Y-276325 (ROCK inhibitor), then treated (before 30min) with TMR labelled 1µM α-SYN protofibrils. Post treatment U-87 MG cells were trypsinized (0.25% trypsin-EDTA solution; Gibco Canada origin #25200-072). Pellet was collected and resuspended in 1X PBS and twice washed with 1X PBS. α-SYN-TMR levels were quantified measuring fluorescence of intensity of TMR at excitation/emission 555/585nm by using flow cytometer (BD LSR II) and analysis was done using analysis software BD LSR-II analysis software.

### Live cell imaging using mitotracker and lysotracker

U-87 MG cells (80,000 cells/well) were seeded on 35 mm glass bottom dish (Cellvis, D35-14-1.5-N) and treated with 1µM TMR labelled α-SYN for 0h, 3h, 6h, 12h and 24 h. The 0h cells were immediately washed before staining with lysotracker and mitotracker. Cells were stained with mitotracker green (Invitrogen #M7514) and lysotracker deep red (Invitrogen #L12492) and were incubated at 37°C for 15 min. Post incubation, the cells were washed with 10% DMEM and time-lapse and z-stack images were taken using confocal microscopy (Zeiss LSM880, Carl Zeiss, Germany). This was done to comprehend the localization of α-SYN and transfer of organelles through TNTs.

### *Live Cell Imaging using* DiD^®^vybrant membrane dye

U-87 MG cells (80,000 cells/well) were seeded on 35 mm dish and treated with 1µM α-SYN as mentioned above. Cells were stained with cell-labelling dye DiD^®^Vybrant (Invitrogen #V22887) to label cell membranes. The dye was diluted with 10% DMEM (phenol red free) in the ratio 1:200 and incubated at 37°C for 20 min. Post incubation, the cells were washed with DMEM for 10 min and images of live cells were taken to visualise the number of thin membrane tubes like TNTs using fluorescence microscope (IX73-Olympus).

### Estimation of cytosolic ROS

2′-7′-Dichlorodifluorescein diacetate (DCFDA) is used to detect ROS production in the cell. U-87 MG cells were seeded in a 24 well plate at a density of 0.1million/well and treated with 1µM α-SYN protofibrils as mentioned above. Post treatment U-87 MG cells were trypsinized (0.25% trypsin-EDTA solution; Gibco Canada origin #25200-072). Pellet was collected and resuspended in DMEM with 10% FBS and 20μM DCFDA. Cells were incubated for 30 min at 37°C then washed with 1X PBS to remove extra dye. ROS levels were quantified measuring fluorescence of DCFDA stained cells at excitation/emission 488/520nm by using flow cytometer (BD LSR II) and analysis was done using analysis software BD LSR-II analysis software.

### Mitochondrial membrane potential using JC-1 dye

JC-1 (5,5,6,6’-tetrachloro-1,1’,3,3’ tetraethylbenzimi-dazolylcarbocyanine iodide) is the indicator of mitochondrial membrane potential. JC-1 is a cationic, lipophilic dye. Normal healthy cells show negative mitochondrial membrane potential where JC-1 dye can enter into the mitochondria and form J-aggregates. The aggregates exhibit excitation/emission maxima at 485/590 nm (red spectrum). Unhealthy cells show higher mitochondrial membrane potential as it loses the balance of electrochemical potential. At this condition, lesser amount of JC-1 dye enters into the mitochondria and retains its monomeric form. These monomers exhibit excitation/emission maxima at 514/529 nm (green spectrum). To know the effect of α-synuclein aggregates on mitochondrial membrane potential, JC-1 assay (BD Biosciences #551302) was done. Astrocytes and U-87 MG cells were seeded at a density of 15,000 cells / well in the glass bottom area of the 35 mm imaging dishes (Cellvis, D35-14-1.5-N). Cells were treated with 1µM α-SYN for 3h-24h. Confocal images were also taken at an excitation of 490 nm and emission at 527 nm (green) and 590 nm (red). The ratio of green fluorescence versus red fluorescence per cell was analysed using Fiji image analysis software to quantify mitochondrial membrane potential.

### Co-culture model and mitochondria transfer

One population of U-87 MG cells were transiently transfected with pLV-mitoDsRed (Addgene #44386; was a gift from Dr. Pantelis Tsoulfas’s lab) plasmid and another population with mEGFP-lifeact-7 (Addgene #58470; was a gift from Michael Davidson) plasmid using lipofectamine 3000 (Invitrogen #44386) transfection reagent. Equal number of transfected cells from both populations were seeded together at a total density of 60,000 cells per glass bottom 35mm dish (Cellvis, #D35-14-1.5-N) and treated with α-SYN protofibrils. After treatment, the cells were fixed with 4 % PFA and images were taken using confocal microscopy (Zeiss LSM880).

### β-galactosidase activity assay

U-87 MG cells were seeded at the density of 10,000 cells per well in a 24 well plate. The cells were treated with 1µM α-SYN for 3h and 24h, these time points were chosen according to the data from other experiments. Post treatment the β-galactosidase activity was measured by using the β-galactosidase staining kit (AKR-100 Cell Biolabs Inc). Brightfield images of the stained cells were taken using colour camera.

### Microscopy

Fluorescence images were taken using Zeiss LSM880 confocal laser scanning microscope (Carl Zeiss, Germany) or fluorescence microscope (IX73-Olympus). The confocal images were taken using objectives Plan-Apochromat 40x/1.40 or 63x/1.40 Oil Dic M27, with the fluorescence filter sets DAPI, FITC, and TRITC (Carl Zeiss, Germany). Sequential images of the different fluorescence channels were taken with 405 nm, 488 nm and 561 nm lasers. The images were captured with pixel dwell of 1.02 µs and each xy-pixel of 220 nm^2^. For all the experiments at least 5-10 images per condition were taken from randomly selected areas. DIC (differential interference contrast) images were captured along with fluorescence channels to understand the morphology and cell boundary for both fixed and live imaging experiments. Time-lapse and z-stacks (6 – 12 stacks of z-scaling ∼ 415 nm) were captured from the bottom to the top of the cells using confocal microscope to identify TNTs and TMs. Trafficking of organelles through TNTs and TMs were tracked from time-lapse images. Wide-field fluorescence microscope (IX73-Olympus) was used to capture images for few experiments using 20X/0.4 NA, and 40X/1.3 NA plan-apochromatic objectives.

### Image analysis

#### a) TNT characterization

Confocal images were analysed using Fiji a Java-based image processing software developed at the National Institutes of Health (NIH) and the Laboratory for Optical and Computational Instrumentation (LOCI). TNTs were characterized by their unique nature to hover and not attach to the substratum, images were taken with z-stacks using confocal microscopy as mentioned above. TNTs (less than 1µM in diameter) were found in the middle z-stacks while TMs are found in the initial z-stacks since they are attached to the surface. Images of z-stacks were reconstructed using 3D volume view plugin of Fiji as described earlier (Dilna et al., 2021). TNTs were manually counted and plotted as the ratio of the number of TNTs by the number of cells per field (Valappil, Raghavan et al. 2022).

#### b) Tracking of organelles

Movements of lysotracker and mitotracker positive vesicles through TNTs and TMs were tracked using the Trackmate (manual) plugin in Fiji. Analysis were done from time lapse videos taken for minimum 30 frames at an interval of 36 seconds for period of 18-20 mins. By setting each organelle as an object we used the semi-automatic tracking method of trackmate plugin. Speed of the organelles were determined from the average displacement between each consecutive image.

#### c) Size analysis

Area of protofibrils, size of aggregates, size of lysosomes and mitochondria were analysed per cell from the randomly taken images of 200-300 cells per condition. The analysis was performed by setting threshold to separate the particles as individual points and then by using ‘Analyze particle’ option of Fiji.

#### d) Intensity analysis

Similarly, expression of Ki67, FAK, pFAK, α-SYN aggregates, lysosomes and ratio of green versus red fluorescence of the JC-1 experiment were analysed by quantifying the intensities of the labelled proteins per cell from the images of 200-300 cells for each condition. Intensities were analysed by drawing and selecting regions of interests using ROI-plugin in Fiji.

#### e) Mitochondria morphology analysis

The quantification of mitochondria branching and branch length were determined using the MiNA plugin in the ImageJ software. We downloaded the MiNA analysis plug in from the Git Hub repository of Stuart lab. We cropped each cell stained with mitochondria and ran the MiNA plugin to determine the mean branch length of mitochondria.

### Statistics

To validate the significance of the analysed data, one-way and two-way ANOVA tests were performed as per the experimental parameters as mentioned in the figure legends. For fig 5C, we performed student t-test since we had only two conditions.

## Author contributions

S.N conceived and conducted the research; A.R, R.K, S.P (JNCASR), R.M (IISc), R.M (JNCASR), S.P (NIMHANS) and S.N designed and interpreted data; A.R, R.K, S.P (JNCASR), S.J, S.C, A.A, M.N.D and M.G performed experiments; A.R, R.K, S.P (JNCASR), S.C, and A.A analysed the data; A.R, R.M (JNCASR) and S.N wrote the paper taking valuable inputs from all the authors.

## Acknowledgement

We thank Mrs. Suma (JNCASR) for the confocal microscopy and Mrs. Usha (JNCASR) for TEM imaging. We thank Dr. Anujith Kumar (MIRM-MAHE) for sharing his resources and Ms Smitha Bhaskar (MIRM-MAHE) for helping in transfection using electroporation technique. We thank Prof. Dipankar Nandi, Indian Institute of Science, for the DCFDA reagent. We thank Dr Lakshmi Balasubramanian (C-CAMP, Bangalore) and Mr. Vedam Pruthvi (MIRM-MAHE) for giving suggestions to develop image analysis flow diagram.

