## Supplementary Material for "Astroglia proliferate upon biogenesis of tunneling nanotubes and clearance of α-synuclein toxicities"

### Supplementary Figures

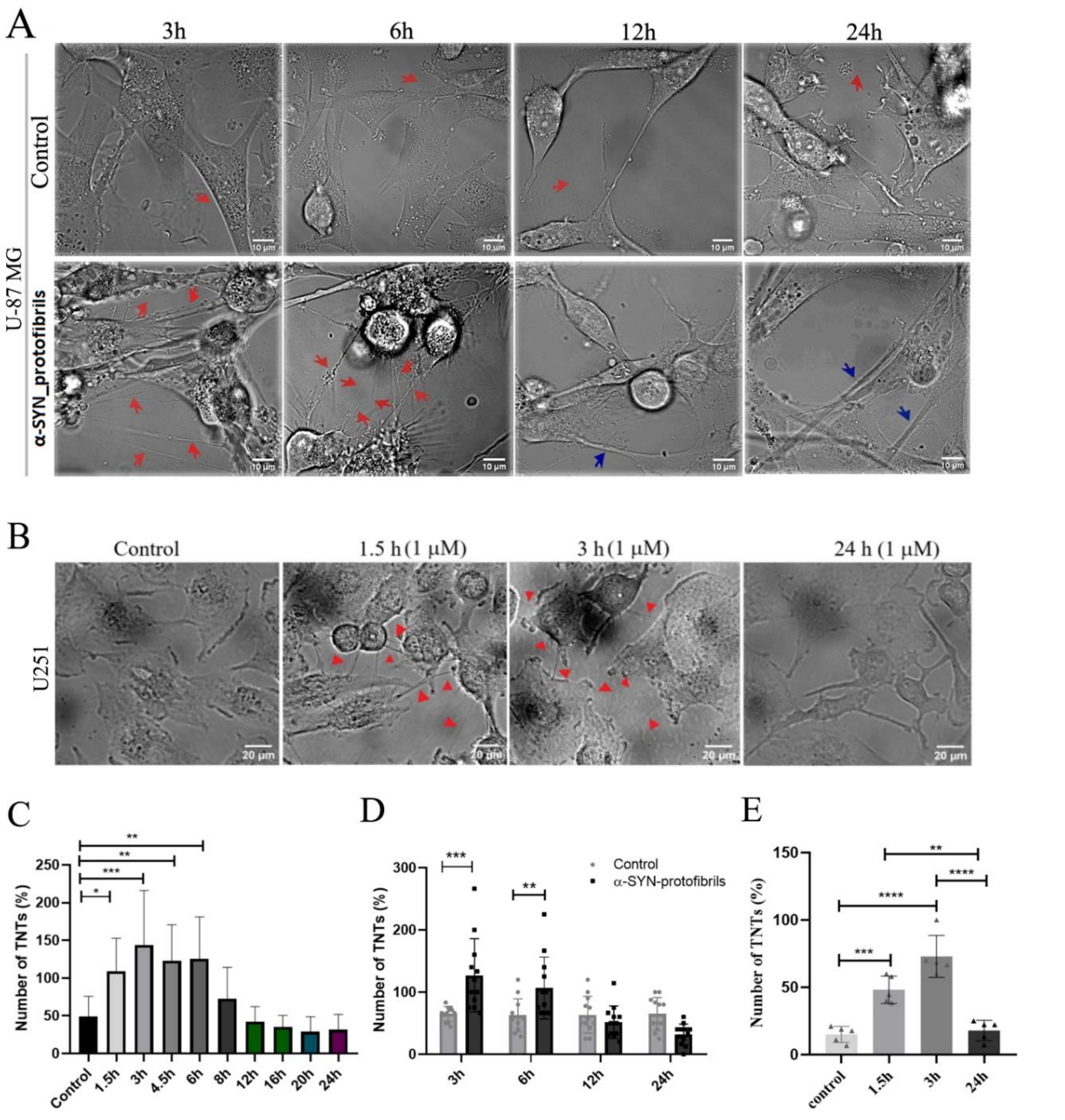


***Figure EV1**: **Transient TNT-biogenesis upon treatment with α-SYN protofibrils in U-87 MG and U-251 cells.*** *A) U-87 MG and B) U251 cells were treated with 1μM α-SYN protofibrils for varying time points from 1.5h to 24h. Increased TNT formation was observed in the early time points 1.5h-3h (marked by red arrow heads) in the DIC images. C) Percentage of TNT numbers were counted across time points varying from 1.5h-24h on treatment with 1μM α-SYN protofibrils in U-87 MG cells. D, E) Percentage of transient TNT-biogenesis were quantified from U-87 MG (D) and U251 (E) cells compared to their respective controls. Scale bars are denoted on the images. Data are expressed as mean ± SD, *** $p \leq 0.001$. Statistics were analysed using one-way ANOVA (EV1C and EV1E) and one-way ANOVA (EV1D). n=3.*

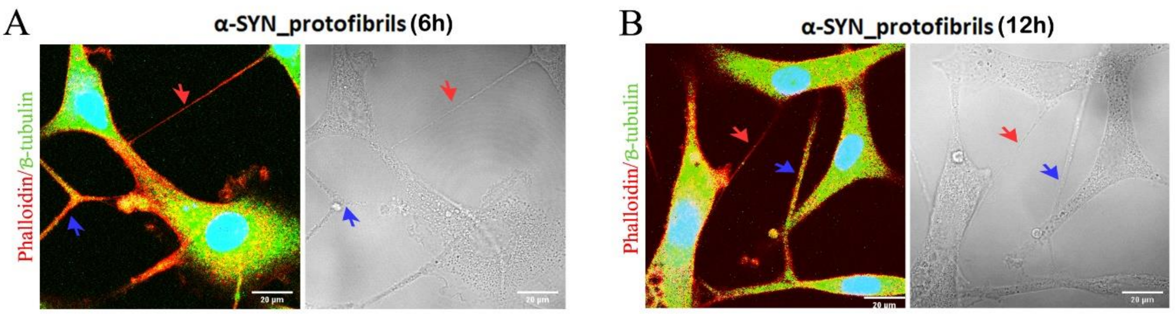


***Figure EV2: Identification and characterization of TNTs and TMs:*** *A-B) U-87 MG cells treated with 1µM α-SYN protofibrils for 6h and 12h, stained for actin and β-tubulin. A) Phalloidin-positive actin-stained thin TNTs and B) β-tubulin positive thicker TMs. Fluorescence images stained with phalloidin and β-tubulin (on the left panels) at 6h and 12h. Red arrows indicate actin positive TNTs and blue arrows indicate β-tubulin positive TMs. Right panels show DIC images of the same. n =3.*

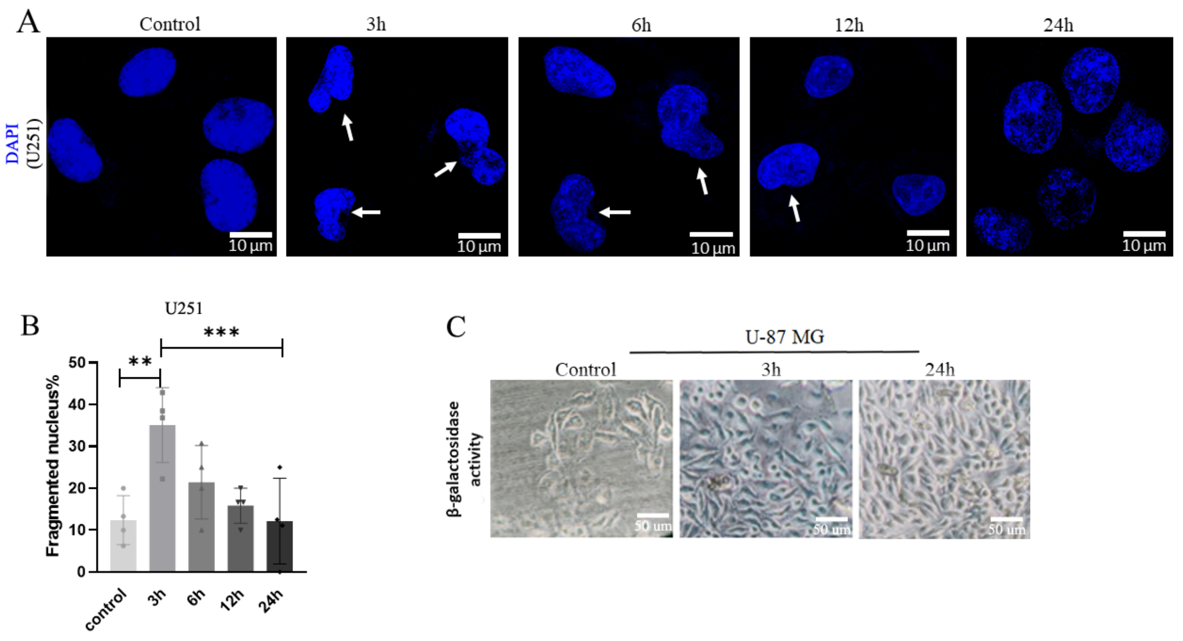


***Figure EV3: α-SYN protofibril induced cellular senescence.*** *A) U251 cells were treated with 1µM α-SYN protofibrils for 3-24h were stained with DAPI. White arrows indicate fragmented nuclei. B) Percentage of the fragmented nucleus per frame were quantified and plotted. Quantifications are done from 5 image frames of a set and each image frame have 10-20 cells. C) U251 cells were treated with 1µM α-SYN protofibrils for 3-24h and checked for β-galactosidase activity. Cellular senescence like characteristics of DNA fragmentation and increased β-galactosidase activity (blue color) was observed at early timepoints. Scale bars are denoted on the images. Data are expressed as mean ± SD, *** $p \leq 0.001$. Statistics were analysed using two-way ANOVA. n=3.*

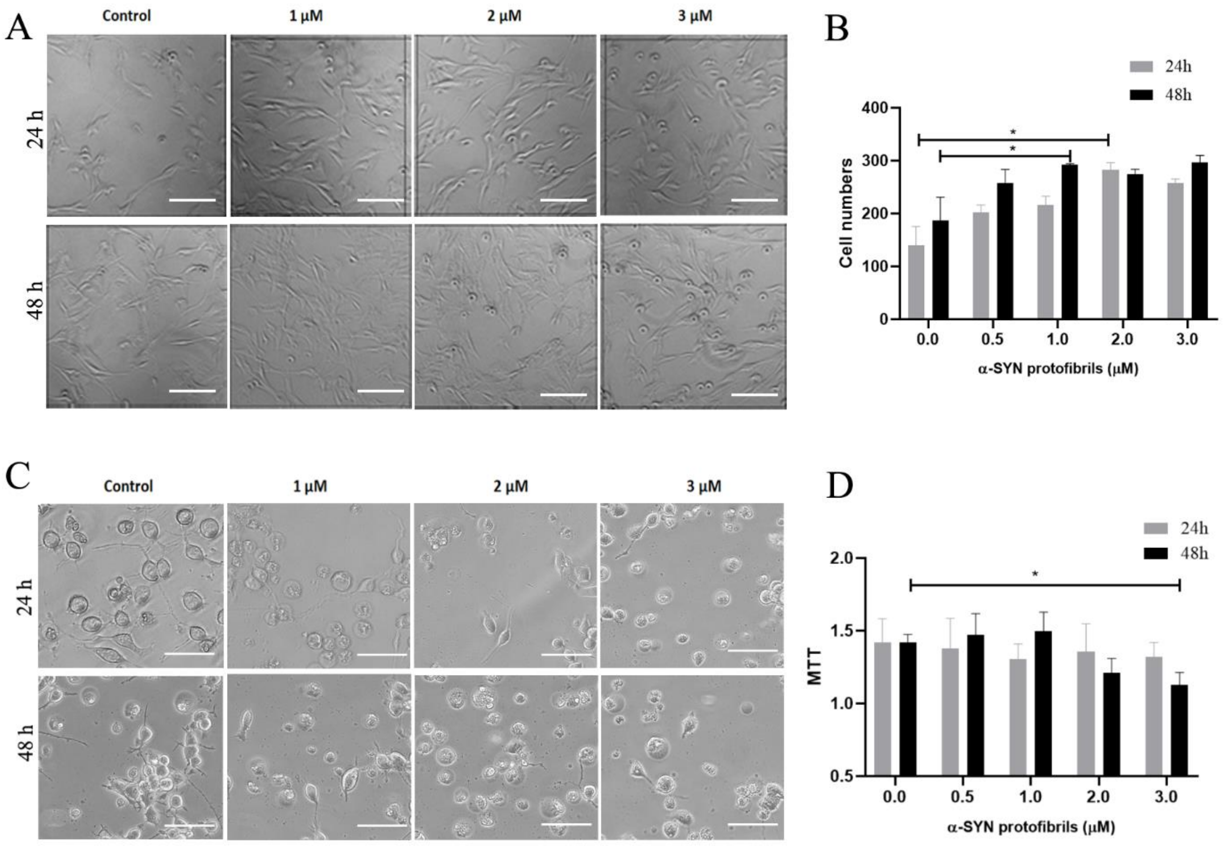


***Figure EV4: α-SYN treatments on U-87 MG and N2a cells in time and concentration dependent manner.*** *A) U-87 MG cells treated with varying concentrations (0.5µM-3µM) of α-SYN protofibrils showed time (24h and 48h) and concentration dependent increase in cell numbers. B) Quantification of the cell numbers. Quantifications are done from 5 image frames of a set and each image frame have 100-250 cells. C) Neurons derived from differentiation of mouse neuroblastoma (N2a) cells treated with toxic α-SYN protofibrils (0.5µM-3µM) for (24h and 48h). The treated cells were imaged and the images show time and concentration dependent occurrence of toxic morphology and floating dead cellsD) MTT assay of N2a cells treated with varying concentrations of α-SYN protofibrils at 24h and 48h. Scale bars are denoted on the images. Data are expressed as mean ± SD, *** $p \leq 0.001$. Statistics were analysed using two-way ANOVA. n=3.*

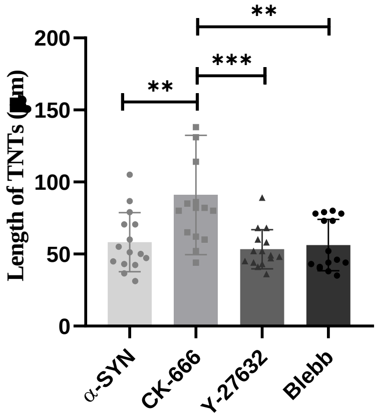


***Figure EV5: Effect of inhibitors on TNT length.*** *Length of TNTs formed in U-87 MG cells after treatment with 1µM α-SYN protofibrils, 50 µM CK-666 (Arp2/3 inhibitor), 5 µM Y-27632 (ROCK inhibitor) and 75 µM Blebbistatin (Blebb) was measured using Image J software. Scale bars are denoted on the images. Quantifications are done from 15 image frames of a set and each image frame have 15-20 cells. Data are expressed as mean ± SD, *** p ≤ 0.001. Statistics were analysed using one-way ANOVA. n=3.*

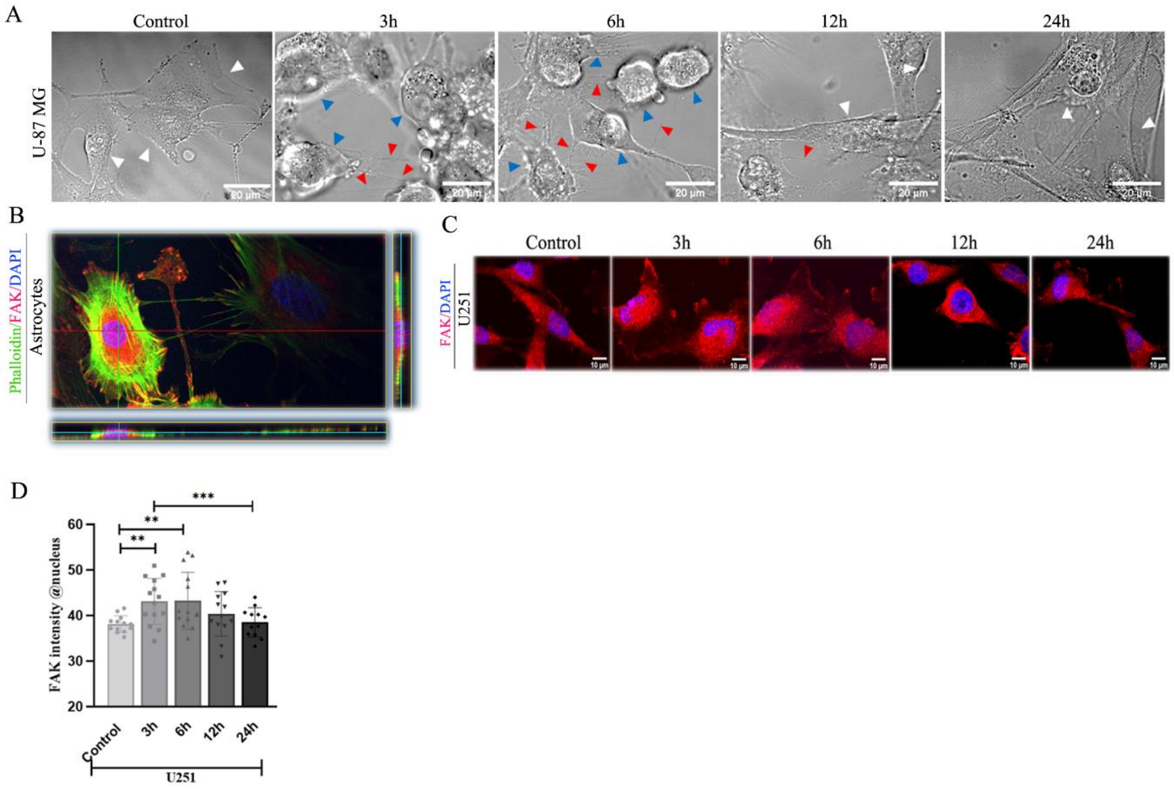


***Figure EV6: Effect of α-SYN on cell adherence and FAK translocation.*** *A) U-87 MG cells were treated with 1µM α-SYN protofibrils for 3-24h, DIC images were taken. Blue arrows indicate non-adherent cells and red arrows indicate TNTs formed from the non-adherent cells. B) Nuclear translocation of FAK was established from 3D view of xz and yz planes. C)Nuclear*

*FAK translocation in U251 cells and D) quantification of the FAK at nucleus. Quantifications are done from 15 image frames of a set and each image frame have 15-20 cells. Scale bars are denoted on the images. Data are expressed as mean ± SD, *** $p \leq 0.001$. Statistics were analysed using two-way ANOVA. n=3.*

#### *Figure legends of supplementary movies:*

***Movie 1:*** *Time-lapse video was captured using DIC channel in confocal microscope with α-SYN protofibrils (1 µM) treated U-87MG cells for 3h. Formation of numerous thin (nanosized in diameter) membrane channels or TNTs between neighbouring cells were observed upon treatment with α-SYN protofibrils for 3h. Organelles were observed to move (indicated by white and black arrows) unidirectional inside the TNTs.*

***Movie 2:*** *Time-lapse video of α-SYN protofibrils (1 µM) treated U-87MG cells for 6h was captured in confocal microscope using DIC channel. Formation of numerous thin (nanosized in diameter) membrane channels or TNTs between neighbouring cells were observed upon α-SYN protofibrils treatment. Organelles were observed to move unidirectionally move (indicated by white arrow) inside the TNTs.*

***Movie 3:*** *Time-lapse video of α-SYN protofibrils (1 µM) treated U-87MG cells for 6h was captured in confocal microscope using fluorescence and DIC channels, to see movements of fluorescently labelled α-SYN-TMR protofibrils and lysotracker positive vesicles inside the TNTs. The composite video shows unidirectional movements of vesicles colocalized (yellow) with α-SYN-TMR (red) and lysotracker (green) in long, thin TNTs (indicated by white arrow).*

***Movie 4:*** *Time-lapse video of α-SYN protofibrils (1 µM) treated U-87MG cells for 3h was captured in confocal microscope using fluorescence and DIC channels, to see movements of mitotracker (cyan) positive vesicles inside the TNTs. The composite video shows unidirectional movements of mitochondria in long, thin TNTs (indicated by white arrow).*

***Movies 5:*** *Time-lapse video captured direct cell-to-cell transfer of α-SYN-TMR (red) accumulated lysotracker (green) positive vesicles (yellow vesicles; indicated by black arrow)* *via TNTs between neighbouring U-87MG cells. The video was captured after 3h of α-SYN protofibril (1 μM) treatments.*

**Movies 6:** *Time-lapse video is able to capture direct cell-to-cell transfer of α-SYN-TMR (red) treated mitotracker positive vesicles (cyan vesicles; indicated by black arrow) via TNTs between neighbouring U-87MG cells. The video was captured after 3h of α-SYN protofibril (1 μM) treatments.*
